# The anti-virus T cell response dominates the anti-cancer response in oncolytic virus therapy

**DOI:** 10.1101/2025.09.22.677905

**Authors:** Meghan J. O’Melia, Kailan Sierra-Davidson, Miranda A. Robert, Lutz Menzel, Lingshan Liu, Pin-Ji Lei, Lance L. Munn, Timothy P. Padera, Sonia Cohen

## Abstract

Oncolytic viruses have strong potential as immunotherapies. By causing cancer cells to die and relieve antigens, these viruses can stimulate robust, systemic immune responses that may eliminate disseminated disease and prevent recurrence. Unfortunately, clinical trials using oncolytic viruses have not induced clearance of metastasis or protection from recurrence. Likewise, the combination of the only FDA-approved oncolytic virus— Talimogene laherparapvec—with immune checkpoint blockade did not improve progression-free or overall survival. Because of these disappointing clinical trials, we sought to measure the ability of oncolytic viruses to induce cancer antigen presentation and the elicitation of cancer antigen-specific immune responses. Our data revealed that despite improved antigen presentation by dendritic cells, priming of cancer antigen-specific T cells was limited. However, viral antigen-specific T cells did develop and were in a phenotypic state to induce an effective response against virally infected cells. These preclinical results were mirrored in human peripheral blood samples. Overall, these data show that oncolytic virus treatment induces a response against the virus itself, but not cancer antigen, explaining the lack of response in metastatic disease. These interesting findings identify a critical mechanism that needs to be overcome to increase the efficacy of oncolytic virus therapy.

## Introduction

Melanoma, the most aggressive form of skin cancer, is a significant clinical problem: approximately 1 in 29-40 people will develop melanoma in their lifetime^1^, with long-term survival rates as low as 35% for later stage disease^1^. In the past decade, immunotherapies have revolutionized the treatment of later stage melanoma due to their potential to treat not only the primary tumor, but also disseminated disease. Further, the development of robust anti-cancer immune responses offer protection from recurrence due to the development of immunological memory ^2–6^. A modern and key immunotherapy that has clinical promise in application to melanoma are oncolytic viruses. These are viruses that preferentially infect cancer cells due to rapid virus proliferation and enhanced ability to interact with tumor cells^7,8^, with several pathogenicity genes removed^9–13^ and potential immunostimulatory genes added^14–16^. The theory behind oncolytic virus therapy is that the virus will kill cancer cells—similar to traditional therapies (e.g. chemotherapy, radiation)— but also induce the release of cancer antigen that will induce local inflammation and a functional, systemic, long-term immune response against cancer antigens^17–19^.

For a cancer antigen-specific response to develop, tumor antigen must first be released from the tumor itself. That antigen must then be taken up, processed, and presented by an antigen-presenting cell (APC), such as a dendritic cell (DC)^20^. An antigen-specific T cell must then interact with the APC to become educated and activated^20^. In theory, oncolytic virus treatment will kill cancer cells, allowing for cancer antigen to be released locally and either be taken up by an APC that migrates to a lymph node (LN) or be transported to a draining LN (dLN) where it is then taken up by an APC (Fig. 1a). In LNs, tissues which concentrate locoregional antigens along with immune cells to allow for more efficient immune cell priming^20^, the APC will present the cancer antigen to cognate lymphocytes to develop an anti-cancer immune response. Despite these theoretical advantages of oncolytic virus therapy, these critical steps (antigen processing, presentation, or development of antigen-specific T cells) have been understudied, both preclinically and clinically, limiting validation of the proposed mechanism of action (Fig. 1a).

**Figure 1:**
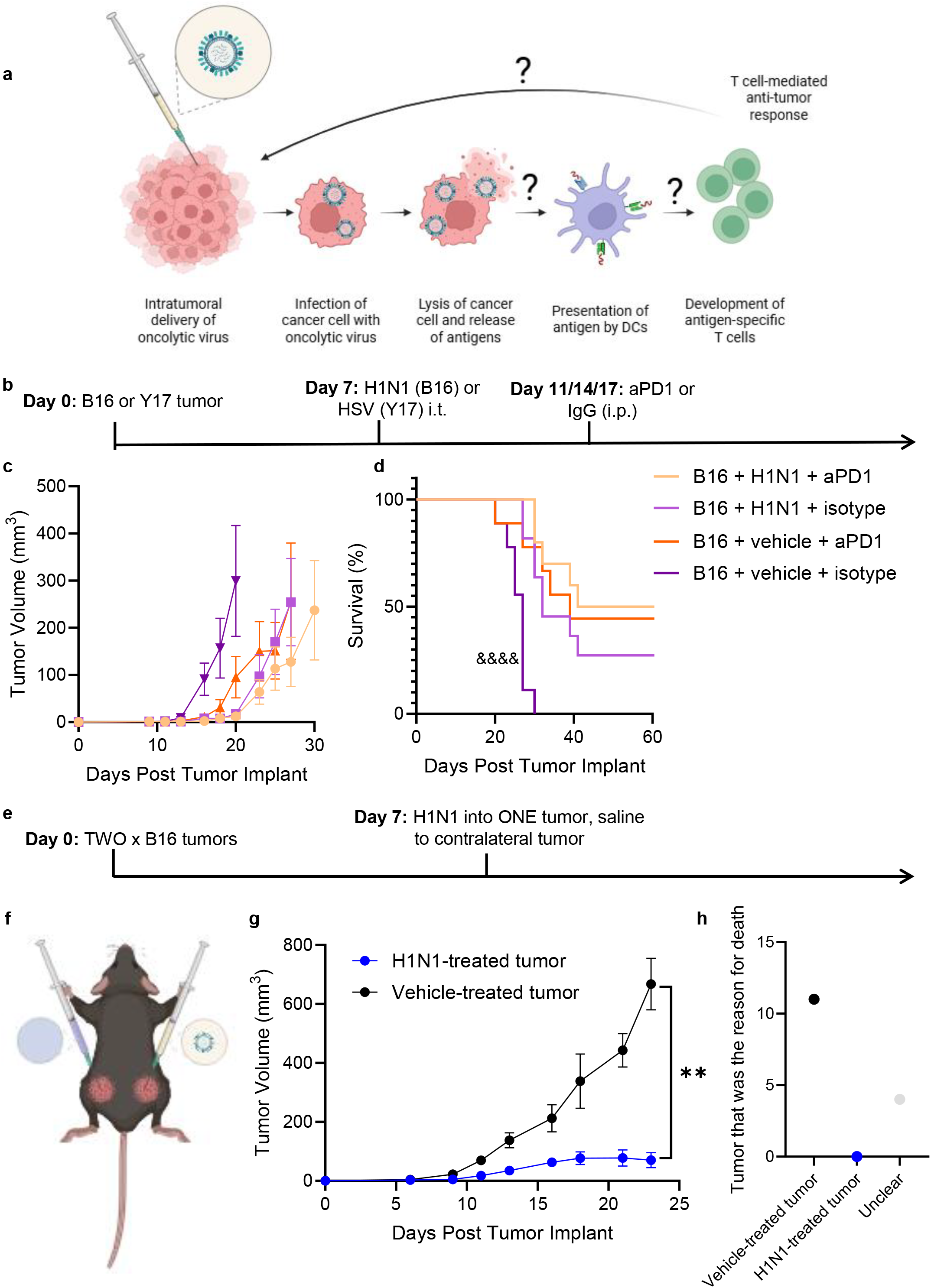
Oncolytic viruses underperform in preclinical models. a, Schematic regarding theory behind the use of oncolytic viruses as melanoma therapies, created in https://BioRender.com. b, Experimental design for combining oncolytic virus with αPD1 therapy in mouse models of melanoma. c, Tumor growth of immunocompetent C57/Bl6 animals bearing B16 tumors treated with H1N1 and/or αPD1 therapy. d, Survival after immunocompetent animals bearing B16 tumors were treated with H1N1 and/or αPD1 therapy. e, Experimental design for analysis of the abscopal effect of oncolytic virus therapy using a two tumor preclinical model. f, Mouse schematic demonstrating tumor location of tumor implantation and treatment (created in https://BioRender.com). g, Tumor growth in two-tumor animals after only one tumor was treated with H1N1 oncolytic virus. h, Reason for sacrifice of two-tumor animals after only one tumor was treated with H1N1 oncolytic virus. * indicates significance by RM-ANOVA; & indicates significance against all others by log-rank test. ** indicates p<0.01, **** indicates p<0.001. n=8-16 animals.

The first and only FDA approved oncolytic virus for use in patients is Talimogene laherparapvec (T-VEC)^21–24^. T-VEC consists of the herpes simplex virus (HSV) with some pathogenic genes removed, and the gene for granulocyte-monocyte colony stimulating factor (GM-CSF) added^25^. In a phase III trial, T-VEC induced an improvement in overall response rate when compared to GM-CSF delivery alone in patients with unresectable stage III melanoma^21^. However, despite an enhancement in survival, only 16.9% of patients achieved complete responses (CR)^21^. As such, T-VEC was combined with anti-programmed death-1 (PD1) monoclonal antibody (mAb) therapy^26^. αPD1 therapy aims to “take the brakes off” the immune system by impairing systems set to tune down an active immune response^27^. The theory behind this clinical trial was that T-VEC-induced anti-tumor antigen-specific T cells would avoid potential exhaustion via αPD1 treatment. However, progression-free or overall survival was not enhanced by this combination therapy when compared to αPD1 alone in a phase III clinical trial^26^.

These disappointing clinical trial results, along with the lack of preclinical knowledge on the mechanisms behind the response to oncolytic viruses, motivated our current study. Here, we measured the major steps required for an effective CD8^+^ T cell response to oncolytic virus therapy, including antigen processing and presentation, as well as the development of a cancer or viral antigen-specific T cell response. We utilized mouse models of melanoma along with human peripheral blood lymphocytes (PBL) before and after melanoma patients were treated with T-VEC to evaluate these critical immunological steps. By testing the theoretical paradigm of oncolytic virus therapy, we identify critical steps that could improve this type of therapy.

## Materials and Methods

### Sex as a Biological Variable

Both male and female animals were used in this study, in approximately equal proportions. Additionally, melanoma cells derived from male mice (YUMMER1.7 [Y17]) and female mice (B16F10 [B16]) are used in this study. The data are presented as combined cohorts as no statistical differences were detected based on sex.

### Animal and Tumor Models

We used 2 different syngeneic melanoma cell lines in this work. B16 melanoma cells (ATCC) or B16 cells modified to present SIINFEKL (B16-OVA) were grown in vitro in Dulbecco’s Modified Eagle’s Medium (DMEM, Gibco, Invitrogen Life Technologies) containing 10% fetal bovine serum (FBS, Atlanta Biologicals) and 1% penicillin-streptomycin (PS, Gibco, Invitrogen Life Technologies). Y17 cells (kind gift of Jessalyn Ubellacker, Harvard Chan School of Public Health) were cultured in DMEM/F-12 medium (Gibco, Invitrogen Life Technologies) supplemented with 10% FBS, 1% PS, and 1X non-essential amino acids (Gibco, Invitrogen Life Technologies). Cells were maintained in a 5% CO2, humidified incubator at 37° C. All cells were recently authenticated and mycoplasma free. B16 or B16-OVA cells (0.25-0.5 × 10^6^) or Y17 cells (10^6^) were prepared to a total volume of 30µL within saline were implanted intradermally in the flank skin of mixed sex (approximately even split) immunocompetent C57/Bl6 mice (aged 5-9 weeks). Animals were monitored every 1-7 days throughout tumor development and progression. All procedures were performed according to the guidelines of the Institutional Animal Care and Use Committee of the Massachusetts General Hospital (Protocol #2011N000085).

### Therapeutic Interventions

H1N1-PR8 virus (Influenza A virus (H1N1) A/Puerto Rico/8/34 VR-95, ATCC) or HSV (kind gift of David Knipe)^28^ were prepared in a total volume of 30µL within saline. This was then injected into the center of the tumor. For combination experiments with checkpoint blockade, αPD1 or α-programmed death ligand-1 (PD-L1) mAbs (BioXCell) were prepared at a concentration of 100ug per 100µL total volume within saline and injected intraperitoneally.

### Enzyme-Linked Immunosorbence Assay (ELISA)

Serum was collected for antibody ELISAs. In short, blood was taken from animals at sacrifice into 1.5mL uncoated Eppendorf tubes. Blood was allowed to clot at room temperature for at least 30 minutes and then samples spun down at 1,000g for 20 minutes. Serum was then removed and maintained at −80° C. To prepare ELISAs, Nunc MaxiSorp high-affinity plates (Thermo Fisher, Inc.) were incubated with ovalbumin (OVA) [10ug OVA (Sigma Aldrich) in 100µL phosphate-buffered saline (PBS)], virus (H1N1) [1µL in 100µL PBS], or 100µL PBS alone overnight. Plates were then washed with 100µL PBS with 1% Tween 20 three times. Serum (5µL serum in 95µL PBS) samples were then added to plates and allowed to incubate for 2 hours at room temperature. Plates were then washed with 100µL PBS three times. Streptavidin conjugated anti-mouse IgG or IgM antibodies (1:1000, Southern Biotech) were then added to plates and allowed to incubate for 2 hours at room temperature. Plates were then washed with 100µL PBS three times. Biotinylated horseradish peroxidase (100µL, R&D Biosystems) was then added to each sample and allowed to incubate for approximately 20 min at room temperature. Stop solution (2N sulfuric acid) was then added to each sample and absorbance read at 450 nm and 540 nm using a multiplex plate reader (MesoScale Discovery).

### Flow Cytometry

Animals were sacrificed in accordance with protocol guidelines. Tissues were resected and maintained on ice in Hank’s Buffered Salt Solution (HBSS). Tissues were then processed by mechanical dissociation through a 70um cell strainer and washed with HBSS. For spleen samples, cells were resuspended in ACK lysis buffer (Thermo Fisher) for 7 min at room temperature and then quenched with approximately 30mL PBS. Samples were centrifuged for 5 minutes at 350g and plated in a 96-well U-bottom plate. The plate was centrifuged for 5 minutes at 350g and Zombie Live/Dead solution (1:100, color in Table 1, Biolegend) was added to samples and allowed to incubate for 30 min at room temperature in the dark. Samples were centrifuged for 5 min at 350g and washed with PBS, then centrifuged again. Fc blocking antibody (1:200, Tonbo Biosciences) was added to samples and allowed to incubate for 20 minutes at room temperature. Samples were centrifuged for 5 min at 350g and surface stain antibodies (Supplemental Table 1) were added. Samples were allowed to incubate for 30 minutes on ice in the dark. Samples were then centrifuged for 5 min at 350g and washed with PBS, and centrifuged again (350g for 5 min). For experiments without intracellular stains, fixative/permeabilization buffer (eBioscience) was added to each sample and allowed to incubate for 30 min on ice in the dark. FACS buffer (1% FBS in PBS) was added to each sample. Samples were then centrifuged for 5 min at 350g, washed with FACS buffer, and centrifuged for 5 minutes at 350g, then maintained in FACS buffer at 4° C in the dark until analyzed via a customized Cytek Aurora flow cytometer. For experiments with intracellular stains, FoxP3 intracellular fixative/permeabilization buffer (eBioscience) was added to each sample and allowed to incubate for 30 min on ice in the dark. Samples were then washed with fixative/permeabilization buffer (eBioscience), centrifuged for 5 min at 350g and intracellular antibodies (Supplemental Table 1) were added and allowed to incubate for 30 min on ice in the dark. Samples were then centrifuged for 5 min at 350g, washed with fixative/permeabilization buffer, centrifuged for 5 min at 350g, and maintained in FACS buffer until analyzed via a customized Cytek Aurora. Data were then analyzed using FlowJo version 10.

### Statistical Analysis

Data are represented as the mean accompanied by standard error of the mean (SEM), and statistics (one-way, two-way, or repeated measures (RM-) analysis of variance (ANOVA) with Tukey’s post-hoc test for grouped analyses; log-rank test for survival analyses; or t-tests, as indicated in figure legends) were calculated using Prism 10 (GraphPad Software Inc., La Jolla, CA, USA). Statistical significance was defined as p<0.05, 0.01, 0.005, 0.001 unless otherwise specified.

### Study Approvals

All experiments were done in accordance with Massachusetts General Hospital Institute for Animal Care and Use Committee policy, per protocol 2011N000085; and Institutional Review Board protocol number DFHCC 11-181.

### Data Availability

All data are presented within the manuscript, and related data are available upon request.

## Results

### Combination of oncolytic viruses with immune checkpoint blockade does not induce synergy, nor do oncolytic viruses induce an abscopal effect

To evaluate the impacts of oncolytic viruses on melanoma tumor growth in preclinical models, we implanted immunocompetent C57/Bl6 animals with B16 or Y17 mouse melanoma lines (Fig. 1b). We then intratumorally injected the PR8 strain of H1N1 influenza (because B16 cells lacked nectin-1, the receptor for HSV/T-VEC (Supplemental Fig. 1) or HSV (the virus that T-VEC is derived from – used in Y17 animals) on day 7 of tumor growth, followed by three doses of αPD1 beginning on day 11 of tumor growth (Fig. 1b). This revealed a slight slowing of tumor growth in all treated groups compared with virus-vehicle combined with isotype IgG control-treated animals (Fig. 1c, Supplemental Fig 2-3).

However, there was no significant difference in tumor growth kinetics between any treatment condition (Fig. 1c, Supplemental Fig. 2-3), implying a lack of synergy between these two treatment subtypes. However, there were no survival differences between any treated group (Fig. 1d, Supplemental Fig. 2-3). These preclinical data replicate what has been shown in clinical trials wherein oncolytic viruses induce a slight increase in overall survival^21^, but there is no benefit to adding αPD1 therapy^26^.

We next investigated the potential of oncolytic viruses to exert an abscopal effect—an anti-cancer response in a lesion not directly targeted by therapy in patients with disseminated disease—within the preclinical setting, as clinical data indicates that oncolytic viruses do not induce this abscopal effect^29^. To investigate potential abscopal effects, we implanted two B16 tumors into immunocompetent C57/Bl6 mice on opposing sides of the flank skin.

One tumor was treated with H1N1 virus, while the other tumor was treated with vehicle (Fig. 1e-f). In this experiment, only the H1N1-treated tumor showed slowing of tumor growth (Fig. 1g, Supplemental Fig. 4). No animals were sacrificed due to complications with the H1N1-treated tumor or the virus itself, only due to either the vehicle-treated tumor or systemic issues due to typical cancer progression (ie. Weight loss) (Fig. 1h). Overall, these data again resemble clinical data in which oncolytic viruses do not exert an abscopal effect^29^. As a whole, these data indicate that preclinical models resemble what has been found in the clinical setting wherein oncolytic viruses do not synergize with αPD1 therapy or induce an abscopal effect on systemic disease.

### Utilizing immunocompromised versus immunocompetent mice with models of melanoma identifies T cells as key players in oncolytic virus responses

To evaluate the role of immune cells in oncolytic virus responses, we analyzed responses in immunodeficient mice compared to immunocompetent animals. First, we compared Nod-scid gamma (NSG) mice, which do not have B, T, or natural killer (NK) cells, compared to immunocompetent C57/Bl6 animals (Fig. 2a). Animals treated with H1N1 on day 7 after B16 implantation revealing that, as expected, tumors developing in immunocompetent animals grew slower when compared to those developing in NSG animals (Fig. 2b, Supplemental Fig. 5). When NSG animals were treated with H1N1, there was no difference in either tumor growth or survival (Fig. 2b-c, Supplemental Fig. 5). However, immunocompetent animals treated with H1N1 exhibited substantial slowing of tumor growth and prolonged survival (Fig. 2b-c, Supplemental Fig. 5). This indicates that T, B, or NK cells are responsible for locoregional impacts of oncolytic viruses in this model. We next evaluated the effects of the virus in Rag1^−/−^ animals, which lack functional B or T cells (Fig. 2a). We saw no difference in either tumor growth or survival after H1N1 treatment in these animals (Fig. 2d-e, Supplemental Fig. 5), indicating that either B or T cells were responsible for the differences in tumor growth seen in Fig. 2b-c. Finally, to evaluate functional antibody generation impacts of B cells, we measured both αH1N1 and αSIINFEKL (tumor antigen)-specific antibodies in serum of immunocompetent animals treated with H1N1 versus vehicle (Fig. 2f), which showed no difference in generation of IgG or IgM antigen-specific antibodies (Fig. 2g-h), indicating no functional antibody generation by B cells. This is not an exhaustive experiment in terms of all that B cells are capable of, but does provide evidence supporting the idea that B cells are not completely responsible for the phenomena we see. As a whole, these data indicate a role for T cells in locoregional effects on tumor growth and survival, which could potentially be exploited to improve long-term responses and development of abscopal effects of oncolytic virus therapy.

**Figure 2:**
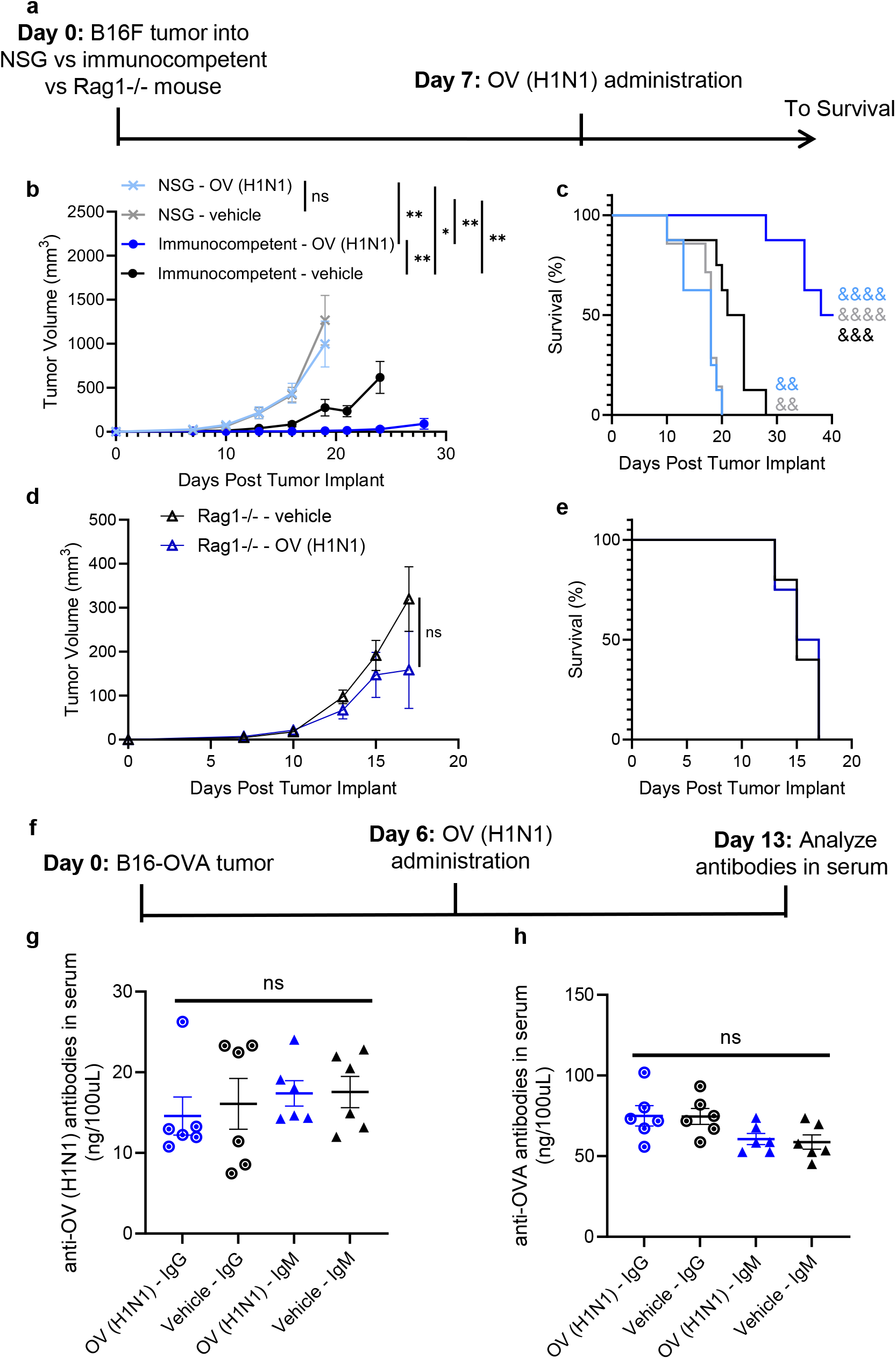
T cells play a dominant role in oncolytic virus response in murine melanoma. a, Experimental design for testing of oncolytic virus in immunocompromised versus immunocompetent animals. b, Tumor growth in NSG versus immunocompetent C57/Bl6 animals after B16 tumor implantation and treatment with H1N1 oncolytic virus. c, Animal survival after NSG versus immunocompetent C57/Bl6 animals were implanted with B16 tumors and treated with H1N1 oncolytic virus. d, Tumor growth after Rag1^−/−^ animals were implanted with B16 tumors and treated with H1N1 virus. e, Survival after Rag1^−/−^ animals were implanted with B16 tumors and treated with H1N1 virus. f, Experimental design for analysis of IgG and IgM development. g, αvirus (H1N1) IgG and IgM within serum of animals treated with H1N1 versus vehicle (saline). h, αOVA IgG and IgM within serum of animals treated with H1N1 versus vehicle (saline). n=6-8 animals. * indicates significance by RM-ANOVA, & indicates significance by log-rank test against all other groups. * indicates p<0.05, ** or && indicate p<0.01, &&& indicates p<0.005, **** or &&&& indicate p<0.001, ns indicates not significant, n=6-8 animals.

### Tumor antigen presentation is enhanced by delivery of an oncolytic virus

As APCs are needed to educate T cells against the tumor, to evaluate the potential of DCs to present tumor antigen in the context of oncolytic virus treatment, we implanted immunocompetent C57/Bl6 animals with B16-OVA tumors (Fig. 3a). Animals were then treated with H1N1 and SIINFEKL (tumor) antigen presentation assessed 1 day later (to evaluate theoretical peak of antigen presentation in the tumor^30^), 4 days later (to evaluate theoretical peak of antigen presentation in the LN^30^), and 19 days later (to evaluate long-term effects of virus) (Fig. 3a) using the 25D1.16 mAb which stains for SIINFEKL within the class I major histocompatibility complex (MHCI). Both the frequency and number of MCHI:SIINFEKL^+^ DCs were enhanced in secondary lymphoid organs (both dLN and non-draining LNs (NdLN), and the spleen) four days after delivery of H1N1 virus when compared to vehicle delivery (Fig. 3b-d, Supplemental Fig. 6). On day 1, antigen presentation was enhanced in the tumor when virus was delivered compared to when vehicle (saline) was delivered, and effects were lost by day 19 (Supplemental Fig. 6-7). Overall, delivery of virus substantially enhanced tumor antigen presentation.

**Figure 3:**
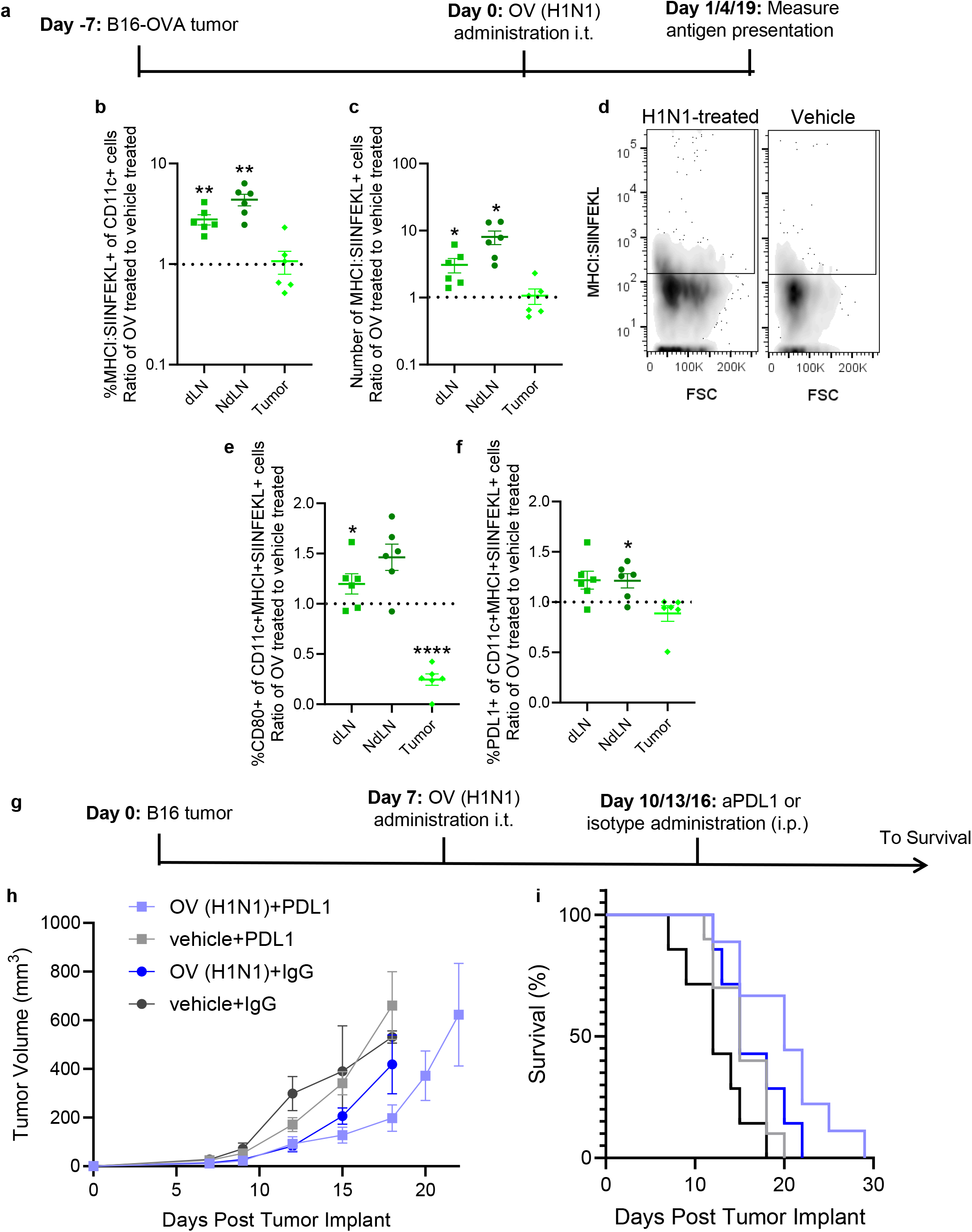
Administration of oncolytic viruses enhances antigen presentation in lymphoid tissues. a, Experimental design for analysis of antigen presentation and DC phenotype in oncolytic virus-treated animals. b, Frequency of DCs presenting tumor antigen (SIINFEKL) within the MHCI in oncolytic virus-treated animals compared to vehicle-treated animals c, number of DCs presenting tumor antigen (SIINFEKL) within the MHCI in oncolytic virus-treated animals compared to vehicle-treated animals. d, representative flow cytometry plots day 4 post-virus administration. e, Frequency of tumor antigen-presenting DCs expressing CD80 in oncolytic virus-treated animals relative to vehicle-treated animals on day 4 post-virus administration. f, Frequency of tumor antigen-presenting DCs expressing PD-L1 in oncolytic virus-treated animals relative to vehicle-treated animals on day 4 post-virus administration. g, Experimental design for evaluation of therapeutic combination of oncolytic viruses with αPD-L1 mAb. h, Tumor growth after animals were treated with oncolytic virus and αPD-L1 mAbs. I, survival after animals were treated with oncolytic virus and αPD-L1 mAbs * indicates significance by t-test against vehicle controls. * indicates p<0.05, ** indicates p<0.01, **** indicates p<0.001, n=6-10 animals.

We next examined the phenotype of DCs presenting tumor antigen, which revealed that within LNs, frequency of CD80^+^ cells were enhanced (Fig. 3e). However, frequency of CD80^+^ cells was decreased in the tumor (Fig. 3e). Tumor-antigen-presenting DCs also had increased PD-L1(Fig. 3e, Supplemental Fig 8). We thus hypothesized that despite enhanced antigen presentation as a whole, DCs presenting tumor antigen could be inducing a less efficient T cell phenotype based on PD-L1 expression (Fig. 3f, Supplemental Fig. 8). To test this, we implanted immunocompetent C57/Bl6 animals with B16 tumors, treated them with H1N1 on day 7 of tumor growth followed by 3 doses of αPD-L1 mAb, and measured tumor growth dynamics and survival (Fig. 3g). This revealed no difference in either tumor growth or survival with addition of αPD-L1 therapy (Fig. 3h-i, Supplemental Fig. 9), indicating that despite enhanced PD-L1 in DCs presenting tumor antigen (Fig. 3f, Supplemental Fig 8), αPD-L1 therapy could not boost anti-cancer immune responses after oncolytic virus therapy.

### Tumor antigen-specific T cell responses were impaired in oncolytic virus treated animals

As MHCI-mediated antigen presentation was enhanced by delivery of H1N1, we next evaluated the impacts of H1N1 administration on the elicitation of cancer antigen-specific CD8^+^ T cell response. To do so, we implanted immunocompetent animals with B16-OVA tumors, administered H1N1 on day 7 post tumor implantation, and then evaluated anti-cancer antigen-specific CD8^+^ T cell responses on days 4, 7, and 11 post-virus delivery (Fig. 4a). This revealed that development of SIINFEKL (tumor antigen)-specific T cells was impaired in both the dLN and tumor of H1N1-treated animals compared to vehicle-treated animals (Fig. 4b-c, Supplemental Fig. 10-12). Because SIINFEKL is a synthetic antigen, we also evaluated development of a gp100-specific T cell response, as B16-OVA cells express gp100 in vitro (Supplemental Fig. 1). gp100-antigen-specific T cells (measured using tetramers) had similar responses when compared to the SIINFEKL antigen-specific T cells, in which dLN, NdLN, and tumors had fewer gp100 antigen-specific T cells in H1N1 oncolytic virus treated animals compared to vehicle treated animals on day 11 after virus delivery (Fig. 4d-e, Supplemental Fig. 10-12). Notably, there were no significant differences in the frequency of CD3^+^ cells or CD3^+^CD8^+^ cells in these tissues (Supplemental Fig. 13), indicating that there was no global T cell effect. Overall, despite enhanced antigen presentation with oncolytic virus treatment (Fig. 3), we saw lower levels of cancer antigen-specific CD8^+^ T cells with delivery of oncolytic virus.

**Figure 4:**
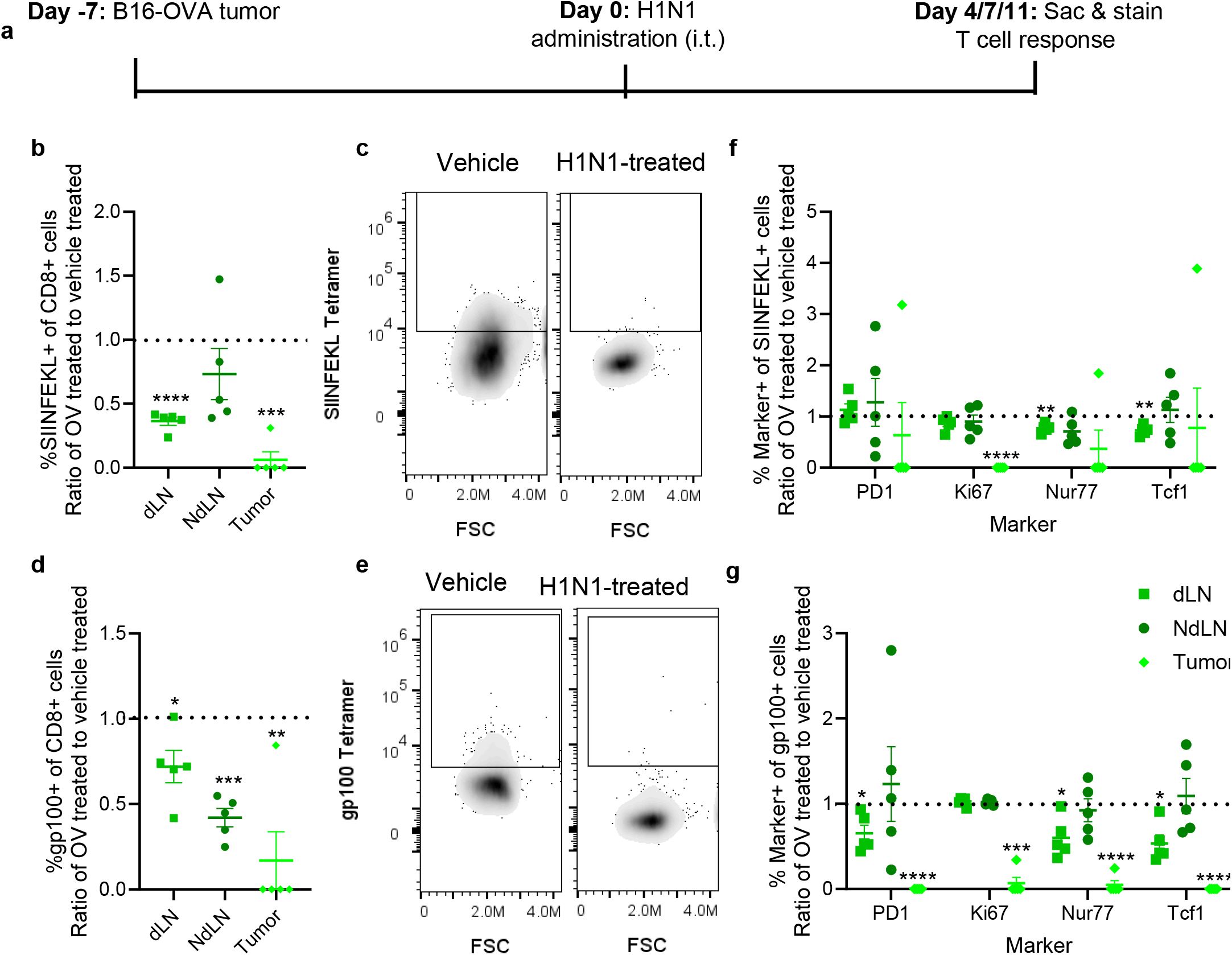
Anti-cancer antigen T cell response is impaired after oncolytic virus treatment. a, Experimental design for analysis of cancer antigen-specific T cells after oncolytic virus delivery. b, Frequency of SIINFEKL antigen-specific T cells in the tumor and LNs of oncolytic virus treated animals relative to vehicle (saline) treated animals on day 11 after virus delivery. c, Representative flow plots for SIINFEKL tetramer staining within the tumor. d, gp100 antigen-specific T cells in the tumor and LNs of oncolytic virus treated animals relative to vehicle (saline) treated animals on day 11 after virus delivery. e, Representative flow plots for gp100 tetramer staining within the tumor on day 11 after virus delivery. f, PD1, Ki67, Nur77, and Tcf1 expression in gp100^+^ CD8^+^ T cells in tumors and lymphoid tissues of oncolytic virus treated animals relative to vehicle (saline) treated animals on day 11 after virus delivery. g, PD1, Ki67, Nur77, and Tcf1 expression in SIINFEKL^+^ CD8^+^ T cells in tumors and lymphoid tissues of oncolytic virus treated animals relative to vehicle (saline) treated animals on day 11 after virus delivery. * indicates significance by t-test relative to vehicle-treated animals. * indicates p<0.05, ** indicates p<0.01, *** indicates p<0.005, **** indicates p<0.001, n=5-6 animals.

We next evaluated the phenotype of the few cancer antigen-specific T cells, as a lower number of higher-quality T cells could potentially be more effective when compared to a higher number of lower-quality cells. Among both the SIINFEKL- and gp100-(tumor antigens) antigen-specific T cells, PD1 (a marker for T cell receptor (TCR) engagement and activation^31^), Ki67 (a marker for proliferation), Nur77 (a marker for very recent TCR engagement^32^), and Tcf1 (a marker for cell stemness^31,33,34^) were all decreased within the tumor when oncolytic virus was delivered compared to vehicle-treated animals (Fig. 4f-g, Supplemental Fig. 12). Taken together, these data indicate that development of functional cancer antigen-specific T cells was impaired by delivery of the virus.

### Viral antigen-specific T cells are present and functional in oncolytic virus treated animals

Our evidence that cancer antigen-specific T cell priming was impaired after oncolytic virus treatment (Fig. 4) was somewhat unexpected, even though our data show that T cells were likely a cause for decreased tumor volume (Fig. 2) and that APC functionality was maintained (Fig. 3) in oncolytic virus treated animals. Thus, we hypothesized that oncolytic virus treatment could be delivering a dominant antigen that dictates CD8^+^ T cell response and limits antigen spreading against cancer antigens. We therefore measured the development of viral antigen-specific T cells in two models of melanoma. To do so, we implanted either B16-OVA or Y17 cells into immunocompetent animals. H1N1 virus (B16-OVA) or HSV (Y17) were delivered intratumorally and viral antigen-specific T cell response measured (Fig. 5a). HSV— similar to T-VEC used clinically^28^—was used to treat the Y17 model as Y17 cells express nectin-1 (Supplemental Figure 1), a known receptor for HSV^35^.

**Figure 5:**
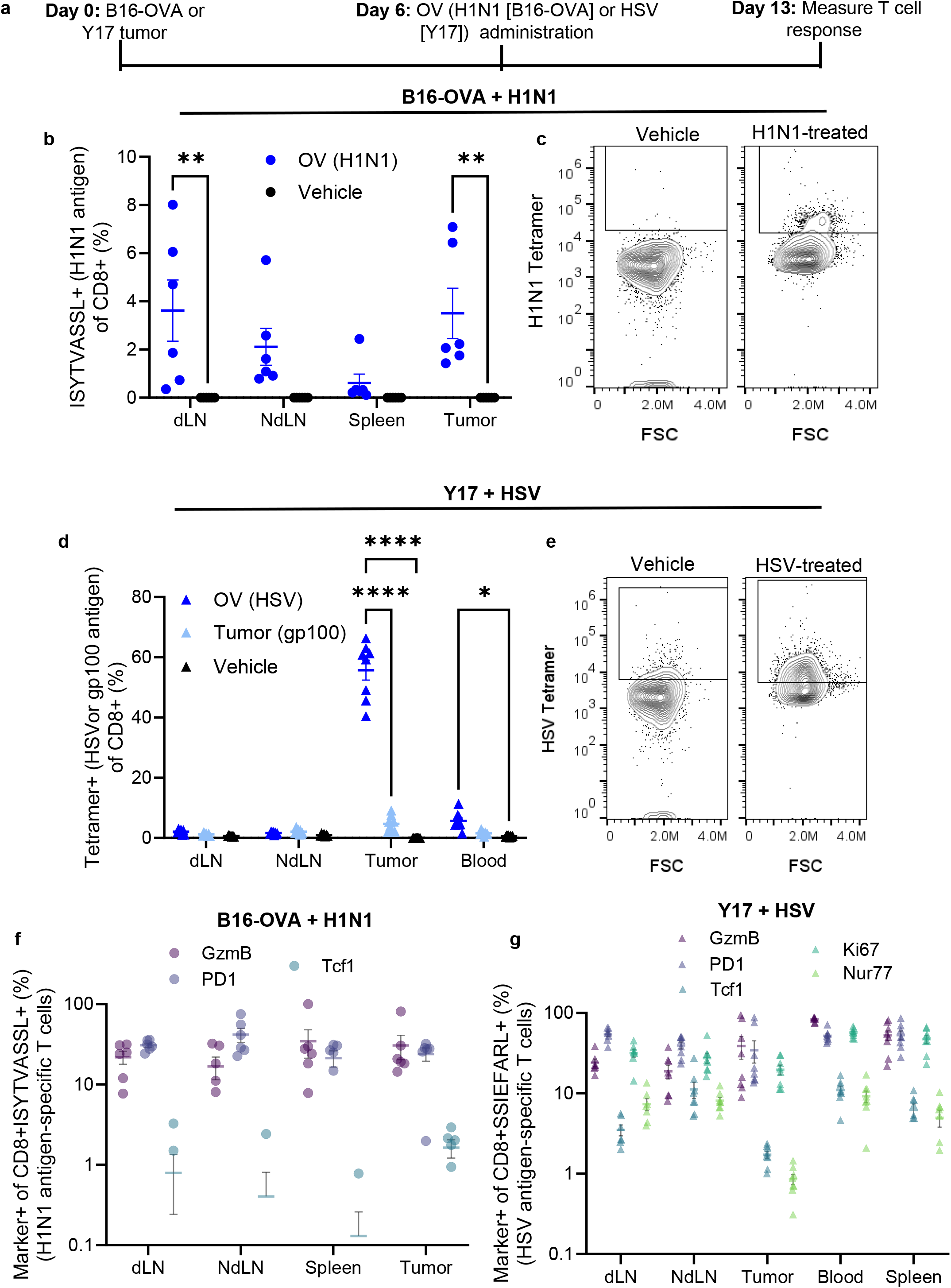
Substantial and effective anti-viral antigen T cell responses develop after oncolytic virus treatment. a, Experimental design for evaluation of anti-viral CD8^+^ T cell responses after oncolytic virus treatment. b, Frequency of CD8^+^ T cells specific for ISYTVASSL (H1N1 antigen) in tissues of interest after B16-OVA-bearing animals were treated with H1N1 virus. c, Representative flow plots of CD8^+^ T cells specific for ISYVASSL (H1N1 antigen) in tumors after B16-OVA-bearing animals were treated with H1N1 virus. d, Frequency of CD8^+^ T cells specific for either HSV or gp100 antigen in tissues of interest after Y17-bearing animals were treated with HSV. e, Representative flow plots of CD8^+^ T cells specific for gp100 antigen in tumors after B16-OVA-bearing animals were treated with H1N1 virus. f, Phenotype of viral antigen-specific T cells in B16-OVA-bearing animals treated with oncolytic virus (no viral antigen-specific T cells were identified in vehicle treated animals). g, Phenotype of viral antigen-specific T cells in Y17-bearing animals treated with oncolytic virus (no viral antigen-specific T cells were identified in vehicle treated animals). * indicates significance by two-way ANOVA. * indicates p<0.05, ** indicates p<0.01, **** indicates p<0.001, n=6-8 animals.

In the B16-OVA model treated with H1N1 virus, we found high frequencies of H1N1 viral antigen-specific T cells in both the dLN and tumor (Fig. 5b-c). In the Y17 model, we found very high levels of HSV antigen-specific T cells within the tumor, and enhanced levels of HSV antigen-specific T cells in the blood (Fig. 5a, d-e). We did not observe any priming of cancer antigen (gp100)-specific T cells in Y17 tumor-bearing animals treated with HSV (Fig. 5d), similar to results we observed in the B16+H1N1 model (Fig. 4).

We next examined the phenotype of these viral antigen-specific T cells. This revealed that among B16-OVA-bearing animals treated with H1N1, there were granzyme B (GzmB) (functionally cytotoxic), PD1 (activated), and Tcf1 (stem-like) viral antigen-specific CD8^+^ T cells (Fig. 5f). Notably, there were no differences in these readouts among total CD8^+^ T cells in these animals (Supplemental Fig. 14). Within the Y17+HSV model, we likewise saw elevated GzmB and PD1 levels as well as Tcf1, Ki67 (proliferating) and Nur77 (recently engaged TCR) positive viral antigen-specific CD8^+^ T cells (Fig. 5g). Overall, this indicates that delivery of either of these two viruses into two different melanoma models induced a functional viral antigen-specific CD8^+^ T cell response, instead of the desired anti-cancer antigen response.

### PBLs from human melanoma patients treated with T-VEC are able to respond to viral but not tumor antigen after ex vivo restimulation

To evaluate whether our preclinical findings were consistent in a clinical context, we utilized human PBLs collected before and after T-VEC delivery from melanoma patients displaying either a CR or no response (NR) to T-VEC. PBLs were thawed, allowed to rest ex vivo for approximately 4 hours, and then stimulated with viral antigen (HSV), cancer antigen (gp100 – gp100 is largely conserved between mice and humans, but human gp100 protein is used in this experiment^36^, or vehicle control (saline) for 6 hours. Our data show that after delivery of T-VEC in a therapeutic setting, CD8^+^ T cells among PBLs stimulated with viral antigen had enhanced GzmB and IL-2 production when compared to cells from patients prior to T-VEC delivery (Fig. 6b-c), implying that these cells were reactive to viral antigen and maintained potentially functional cytotoxicity. Among cells stimulated with cancer antigen (gp100), we saw no response as measured by GzmB and IL-2 production (Fig. 6b-c), implying that these cells were not able to exert a cytotoxic response against cancer antigen. Interestingly, there were no differences between CR and NR patients in these trends (Fig. 6b-c). These data are consistent with our data from preclinical models. We also measured CD45RO (a memory marker, indicating better αPD1 responses), tumor necrosis factor-α (TNFα, an inflammatory cytokine), CD38 (a marker of exhausted resident memory T cells), 4-1BB (a marker of activation), PD1, and CD27 (a marker of naïve and memory[non-effector] cells) in these samples, and did not find any differences among these markers (Supplemental Fig. 15) when comparing pre- and post-T-VEC samples.

**Figure 6:**
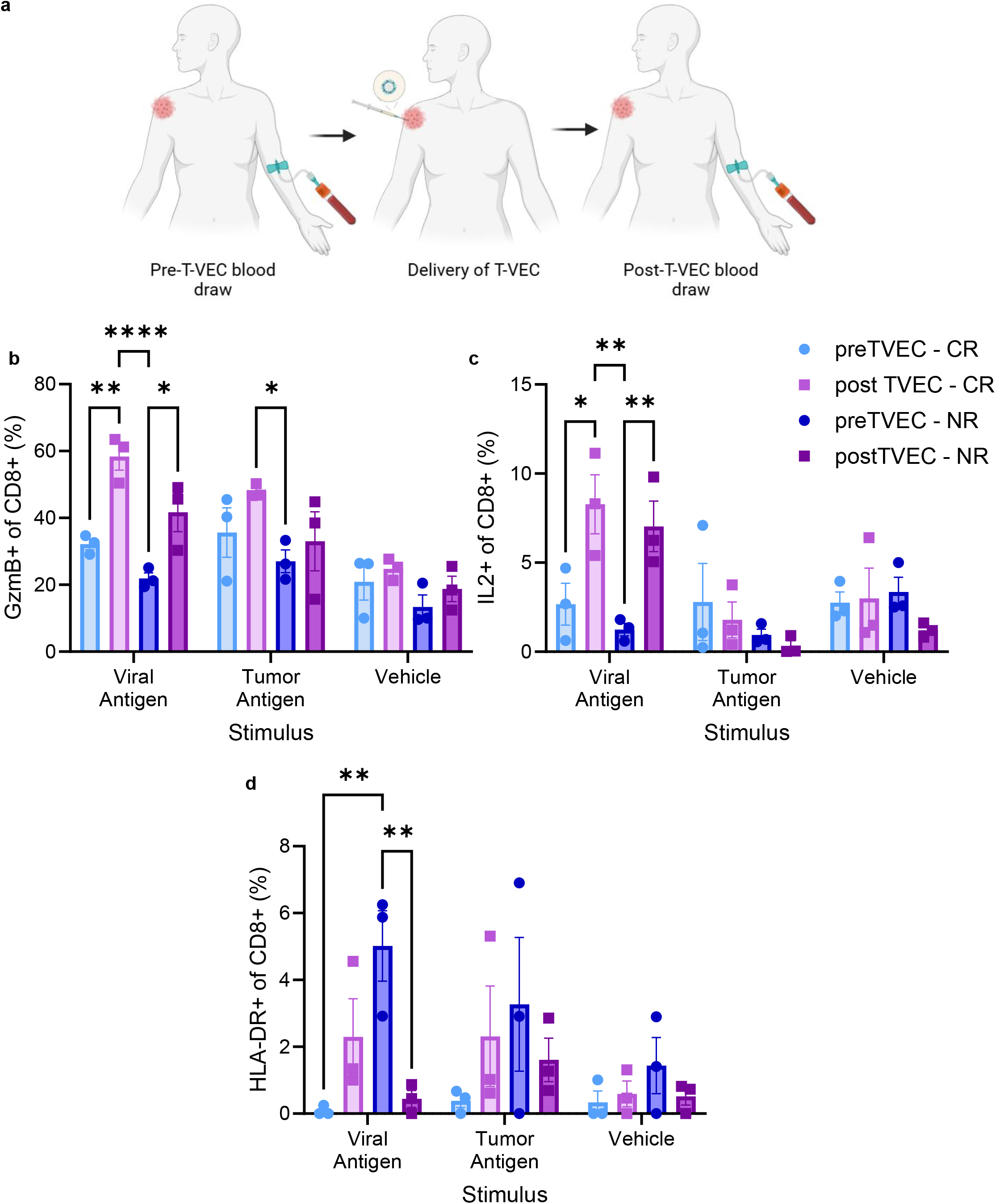
Oncolytic virus delivery induces viral but not cancer antigen-specific T cells in human PBLs. a, Experimental design for evaluation of clinical samples, created in https://BioRender.com. b, Frequency of CD8^+^ cells stained positive for GzmB after ex vivo stimulation with viral antigen, cancer antigen, or vehicle ex vivo. c, Frequency of CD8^+^ cells stained positive for IL2 after ex vivo stimulation with viral antigen, cancer antigen, or vehicle ex vivo. d, Frequency of CD8^+^ cells stained positive for HLA-DR after ex vivo stimulation with viral antigen, cancer antigen, or vehicle ex vivo. * indicates significance by two-way ANOVA with Tukey’s post-hoc comparison. * indicates p<0.05, ** indicates p<0.01, **** indicates p<0.001, n=3 CR, 3 NR (pre and post-T-VEC for each patient). Where significance is not shown, p>0.05.

### HLA-DR may be a biomarker for response to T-VEC therapy clinically

In comparing samples from CR and NR patients, we identified that HLA-DR frequency, a marker indicating late or chronic T cell activation, was higher among NR patients before T-VEC delivery in viral antigen-stimulated T cells when compared to CR patients before T-VEC delivery (Fig. 6d). Similarly, HLA-DR frequency was higher in NR patients prior to T-VEC delivery compared to NR patients after T-VEC delivery (Fig. 6d). These trends were similar (but did not reach significance) in tumor antigen or vehicle-stimulated T cells (Fig. 6d). This should be studied further as a potential biomarker to stratify patients that may be more responsive to T-VEC.

## Discussion

Our results indicate that delivery of oncolytic viruses induces a CD8^+^ T cell effect against only viral antigen instead of cancer antigen. Here, we show this phenomena in multiple preclinical models of melanoma, along with human PBLs from T-VEC-treated patients. The lack of anti-cancer antigen response occurs in spite of enhanced cancer antigen presentation after oncolytic virus delivery. As cancer cells within the tumor that has been injected intratumorally with the oncolytic virus will be infected by the virus, the anti-viral cytotoxic T cell response can explain responses in the primary tumor. Since distant metastatic sites are unlikely to be virally infected, our data also explain the lack of an abscopal effect. Understanding the mechanism behind the lack of antigen spreading or generation of anti-cancer antigen-specific T cell responses after oncolytic virus therapy provides an opportunity to develop next generation oncolytic virus therapy that can lead to robust systemic anti-cancer immune responses.

These data and conclusions have substantial ramifications for the use of oncolytic viruses clinically. Modifications to therapeutic regimens should be considered – including alterations in the oncolytic virus itself to decrease immunogenicity of viral antigens. Further investigation into why the DCs presenting cancer antigens are not able to activate T-cell responses in the face of the strong anti-viral response is also necessary. Understanding these fundamental immunological mechanisms —which may have evolved to prevent auto-immunity after natural exposure to lytic viruses—may also provide insights for next generation oncolytic therapies.

Why the viral antigen-specific T cell response dominates cancer antigen T cell response is an interesting open question beyond just oncolytic virus therapy. For example, in the context of concurrent infections, how does one antigen response dominate the other? In the context of cancer, which antigens are prioritized as targets? Some potential explanations for this phenomena include overloading of antigen and restriction of antigen loading during DC maturation^37–41^; feedback loops such as inhibitory receptor (ie. PD-L1) expression and Treg activation^42–44^; and deletion of tumor-specific T cells in the thymus due to similarities to self antigen^45–47^, all of which should be considered in generation of new generation oncolytic viruses and other immunotherapies. Further investigation into these fundamental immunological questions will lead to exciting new discoveries.

## Acknowledgments

We thank Peigen Huang, Anna Khachatryan, Mark Duquette, and Hengbo Zhou for their hard work and technical assistance in these studies. We additionally thank Jessalyn Ubellacker for the kind gift of the Y17 cells. We thank the NIH Tetramer Core Facility (contract number 75N9302D00005) for providing tetramers.

## Supplemental Figure Legends

**Supplemental Figure 1: Individual tumor growth curves after animals bearing B16 tumors were treated with H1N1 and αPD1**.

**Supplemental Figure 2: Tumor growth and survival of animals bearing Y17 tumors treated with HSV and αPD1**.

**Supplemental Figure 3: Individual tumor growth curves from animals bearing two B16 tumors, one of which was treated with H1N1 virus**.

**Supplemental Figure 4: Individual growth curves after NSG, immunocompetent C57/Bl6, or Rag1^−/−^ animals bearing B16 tumors were treated with H1N1 virus**.

**Supplemental Figure 5: Antigen presentation data within spleen samples**. a, Ratio of each readout (x-axis) within the spleen of H1N1-treated compared to vehicle-treated animals on day 1 post-virus treatment. b, Ratio of each readout (x-axis) within the spleen of H1N1-treated compared to vehicle-treated animals on day 4 post-virus treatment. c, Ratio of each readout (x-axis) within the spleen of H1N1-treated compared to vehicle-treated animals on day 19 post-virus treatment. * indicates significance by t-test (relative to vehicle-treated animals). * indicates p<0.05, *** indicates p<0.005, **** indicates p<0.001, n=6 animals.

**Supplemental Figure 6: Day 1 and day 19 evaluation of APCs**. a, Frequency of MHCI:SIINFEKL^+^ cells of all CD11c^+^ cells in animals treated with H1N1 virus relative to vehicle treated animals on day 1 after delivery of H1N1 virus. b, Number of MHCI:SIINFEKL^+^ cells in animals treated with H1N1 virus relative to vehicle treated animals on day 1 after delivery of H1N1 virus. c, Frequency of CD80^+^ cells of all MHCI:SIINFEKL^+^ cells in animals treated with H1N1 virus relative to vehicle treated animals on day 1 after delivery of H1N1 virus. d, Frequency of PD-L1^+^ cells of all MCHI:SIINFEKL^+^ cells in animals treated with H1N1 virus relative to vehicle treated animals on day 1 after delivery of H1N1 virus. e, Frequency of MHCI:SIINFEKL^+^ cells of all CD11c^+^ cells in animals treated with H1N1 virus relative to vehicle treated animals on day 19 after delivery of H1N1 virus. f, Number of MHCI:SIINFEKL^+^ cells in animals treated with H1N1 virus relative to vehicle treated animals on day 19 after delivery of H1N1 virus. g, Frequency of CD80^+^ cells of all MHCI:SIINFEKL^+^ cells in animals treated with H1N1 virus relative to vehicle treated animals on day 19 after delivery of H1N1 virus. h, Frequency of PD-L1^+^ cells of all MCHI:SIINFEKL^+^ cells in animals treated with H1N1 virus relative to vehicle treated animals on day 19 after delivery of H1N1 virus. * indicates significance by t-test (relative to vehicle-treated animals). * indicates p<0.05, ** indicates p<0.01, *** indicates p<0.005, **** indicates p<0.001, n=6 animals.

**Supplemental Figure 7: Phenotype of total CD11c**^**+**^ **cells**. a, CD80^+^ cells of total (ie. not exclusively MHCI:SIINFEKL^+^) CD11c^+^ cells in H1N1-treated animals relative to vehicle-treated animals on day 1 after virus treatment. b, CD80^+^ cells of total (ie. not exclusively MHCI:SIINFEKL^+^) CD11c^+^ cells in H1N1-treated animals relative to vehicle-treated animals on day 4 after virus treatment. c, CD80^+^ cells of total (ie. not exclusively MHCI:SIINFEKL^+^) CD11c^+^ cells in H1N1-treated animals relative to vehicle-treated animals on day 19 after virus treatment. d, PD-L1^+^ cells of total CD11c^+^ cells in H1N1-treated animals relative to vehicle-treated animals on day 1 after virus treatment. e, PD-L1^+^ cells of total CD11c^+^ cells in H1N1-treated animals relative to vehicle-treated animals on day 4 after virus treatment. f, PD-L1^+^ cells of total CD11c^+^ cells in H1N1-treated animals relative to vehicle-treated animals on day 19 after virus treatment. n=6 animals.

**Supplemental Figure 8: Individual tumor growth curves after B16-bearing animals were treated with H1N1 and αPD-L1 therapy**.

**Supplemental Figure 9: Cancer antigen-specific T cell responses within the spleen**. a, Frequency of either gp100 or SIINFEKL antigen-specific CD8^+^ T cells in the spleen in H1N1-treated animals relative to vehicle-treated animals on day 11 after virus delivery. b, Phenotype of gp100-specific T cells within the spleen in H1N1-treated animals relative to vehicle treated animals on day 11 after virus delivery. c, Phenotype of SIINFEKL-specific T cells within the spleen in H1N1-treated animals relative to vehicle treated animals on day 11 after virus delivery. n=5 animals.

**Supplemental Figure 10: Cancer antigen-specific T cell responses on day 4 or 7 after virus delivery**. a, gp100 antigen-specific CD8^+^ T cells in H1N1-treated animals relative to vehicle-treated animals on day 4 after virus delivery. b, SIINFEKL antigen-specific CD8^+^ T cells in H1N1-treated animals relative to vehicle-treated animals on day 4 after virus delivery. c, gp100 antigen-specific T cells in H1N1-treated animals relative to vehicle-treated animals on day 7 after virus delivery. d, SIINFEKL antigen-specific T cells in H1N1-treated animals relative to vehicle-treated animals on day 7 after virus delivery. n=5 animals.

**Supplemental Figure 11: Nectin-1 and gp100 staining in B16 and Y17 models**. a, Nectin-1 staining in Y17 cells, comparing unstained to stained cells (from culture). b, Nectin-1 staining in B16 cells, comparing unstained to stained cells (from culture). c, gp100 staining in Y17 tumors (in vivo). d, gp100 staining in B16 cells (from culture).

**Supplemental Figure 12: Phenotype of tumor antigen-specific T cells on day 4 or 7 after virus delivery**. a, Phenotypic marker (x-axis) of gp100 specific CD8^+^ T cells in H1N1-treated animals relative to vehicle-treated animals on day 4 after virus delivery. b, Phenotypic marker (x-axis) of SIINFEKL specific CD8^+^ T cells in H1N1-treated animals relative to vehicle-treated animals on day 4 after virus delivery. c, Phenotypic marker of gp100 specific CD8^+^ T cells in H1N1-treated animals relative to vehicle treated animals on day 7 after virus delivery. d, Phenotypic marker of SIINFEKL specific CD8^+^ T cells in H1N1-treated animals relative to vehicle treated animals on day 7 after virus delivery. n=5 animals.

**Supplemental Figure 13: CD3 and CD8 frequencies after viral delivery**. a, Frequency of CD3^+^ of total live cells in H1N1-treated animals relative to vehicle-treated animals on day 4 after delivery of virus. b, Frequency of CD3^+^ of total live cells in H1N1-treated animals relative to vehicle-treated animals on day 7 after delivery of virus. c, Frequency of CD3^+^ of total live cells in H1N1-treated animals relative to vehicle-treated animals on day 11 after delivery of virus. d, Frequency of CD8^+^ of total CD3^+^ cells in H1N1-treated animals relative to vehicle-treated animals on day 4 after delivery of virus. e, Frequency of CD8^+^ of total CD3^+^ cells in H1N1-treated animals relative to vehicle-treated animals on day 7 after delivery of virus. f, Frequency of CD8^+^ of total CD3^+^ cells in H1N1-treated animals relative to vehicle-treated animals on day 11 after delivery of virus. n=5 animals.

**Supplemental Figure 14: Phenotype of total CD8**^**+**^ **T cells after virus delivery**. a,

Frequency of CD8^+^ (total, not necessarily Ag-specific) stained positive for GzmB in B16-OVA-bearing animals treated with H1N1 or vehicle. b, Frequency of CD8^+^ (total, not necessarily Ag-specific) stained positive for PD1 in B16-OVA-bearing animals treated with H1N1 or vehicle. c, Frequency of CD8^+^ (total, not necessarily Ag-specific) stained positive for Tcf1 in B16-OVA-bearing animals treated with H1N1 or vehicle. ns indicates p>0.05, n=5 animals.

**Supplemental Figure 15: Non-significant markers in clinical samples**. a, CD45RO staining in human PBLs stimulated ex vivo with viral antigen, cancer antigen or vehicle. b, TNFα staining in human PBLs stimulated ex vivo with viral antigen, cancer antigen or vehicle. c, CD38 staining in human PBLs stimulated ex vivo with viral antigen, cancer antigen or vehicle. d, 4-1BB staining in human PBLs stimulated ex vivo with viral antigen, cancer antigen or vehicle. e, PD1 staining in human PBLs stimulated ex vivo with viral antigen, cancer antigen or vehicle. f, CD27 staining in human PBLs stimulated ex vivo with viral antigen, cancer antigen or vehicle. n=3 CR, 3 NR.

**Supplemental Table 1:** Flow cytometry antibodies.

| Color | Target | Clone | Vendor | Cat # |
| --- | --- | --- | --- | --- |
| Antigen Presentation analysis |  |  |  |  |
| APC-Cy7 | Live/Dead |  | eBioscience | 65-0865-14 |
| PE | MHCI:SIINFEKL | 25D1.16 | Biolegend | 141604 |
| APC | CD11c | N418 | Biolegend | 117310 |
| FITC | PD-L1 | MIH2 | Biolegend | 393606 |
| PE-Cy7 | CD80 | 16-10A1 | Biolegend | 104734 |
| PerCP-Cy5.5 | CD45 | 30-F11 | Biolegend | 103132 |
| Tumor antigen-specific T cell analysis |  |  |  |  |
| UV | Live/Dead |  | Biolegend | 423108 |
| BV510 | CD45 | S18009F | Biolegend | 157219 |
| PE-Cy7 | CD8 | SK1 | Biolegend | 344712 |
| PerCP | CD3 | 17A2 | Biolegend | 100288 |
| BV785 | PD1 | 29F.1A12 | Biolegend | 135225 |
| BV421 | SIINFEKL (Tetramer) |  | NIH Tetramer Core |  |
| APC | gp100 (Tetramer) |  | NIH Tetramer Core |  |
| BV570 | CD44 | IM7 | Biolegend | 103037 |
| AF594 | Nur77 | 1E10A15 | Biolegend | 653504 |
| PE | Tcf1 | 7F11A10 | Biolegend | 655208 |
| BV650 | Ki67 | 11F6 | Biolegend | 151215 |
| Viral antigen-specific T cell analysis |  |  |  |  |
| BV510 | CD45 | S18009F | Biolegend | 157219 |
| BV421 | ASNENMETM (Tetramer) |  | NIH Tetramer Core |  |
| APC | ISYVASSL (Tetramer) |  | NIH Tetramer Core |  |
| PerCP | CD3 | 17A2 | Biolegend | 100288 |
| PE-Cy7 | CD8 | SK1 | Biolegend | 344712 |
| FITC | CD4 | GK1.5 | Biolegend | 100406 |
| BV785 | PD1 | 29F.1A12 | Biolegend | 135225 |
| PE | Tcf1 | 7F11A10 | Biolegend | 655208 |
| PE-Cy5 | GzmB | QA16A02 | Biolegend | 372226 |
| PerCP-Cy5.5 | FoxP3 | FJK-16s | eBioscience | 45-5773-82 |
| UV | Live/Dead |  | Biolegend | 423108 |
| nectin 1 staining |  |  |  |  |
| PE | nectin-1 | R1.302 | Thermo | MA5-28593 |
| T cell responses in Y17 model |  |  |  |  |
| BV421 | HSV (Tetramer) |  | NIH Tetramer Core |  |
| APC | gp100 (Tetramer) |  | NIH Tetramer Core |  |
| PerCP | CD3 | 17A2 | Biolegend | 100288 |
| PE-Cy7 | CD8 | SK1 | Biolegend | 344712 |
| FITC | GzmB | QA18A28 | Biolegend | 396404 |
| BV785 | PD1 | 29F.1A12 | Biolegend | 135225 |
| PE | Tcf1 | 7F11A10 | Biolegend | 655208 |
| BV570 | CD44 | IM7 | Biolegend | 103037 |
| BV650 | Ki67 | 11F6 | Biolegend | 151215 |
| AF594 | Nur77 | 1E10A15 | Biolegend | 653504 |
| BV510 | CD45 | S18009F | Biolegend | 157219 |
| NIR | Live/Dead |  | Biolegend | 423106 |
| T cell responses in human PBLs |  |  |  |  |
| PerCP-Cy5.5 | CD45 | HI30 | Biolegend | 304028 |
| AF488 | CD3 | HIT3a | Biolegend | 300320 |
| PE-Cy7 | CD8 | SK1 | Biolegend | 344712 |
| AF700 | CD4 | SK3 | Biolegend | 344622 |
| BV421 | PD1 | NAT105 | Biolegend | 367422 |
| APC | GzmB | QA16A02 | Biolegend | 372204 |
| PE | IL2 | MQ1-17H12 | Biolegend | 500307 |
| BV510 | Ki67 | 11F6 | Biolegend | 151225 |
| BV711 | TNFα | MAb11 | Biolegend | 502940 |
| BV650 | FoxP3 | 206D | Biolegend | 320134 |
| BV785 | 4-1BB | 4B4-1 | Biolegend | 309850 |
| APC-Cy7 | CD38 | S17015A | Biolegend | 397118 |
| BV570 | HLA-DR | L243 | Biolegend | 307638 |
| PE-Cy5 | CD27 | O323 | Biolegend | 302858 |
| AF594 | CD45RA | HI100 | Biolegend | 304160 |
| BV605 | CD45RO | UCHL1 | Biolegend | 304238 |
| NIR | Live/Dead |  | Biolegend | 423106 |

## Abbreviations

ANOVA: analysis of variance
APC: Antigen-presenting cell
B16: B16F10
CR: complete response
DC: dendritic cell
dLN: draining lymph node
DMEM: Dulbecco’s Modified Eagle’s Medium
ELISA: enzyme-linked immunosorbence assay
FBS: fetal bovine serum
GM-CSF: granulocyte-monocyte colony stimulating factor
GzmB: granzyme B
HBSS: Hank’s buffered salt solution
HSV: herpes simplex virus
LN: lymph node
mAb: monoclonal antibody
MHCI: class I major histocompatibility complex
NdLN: non-draining lymph nodes
NR: no response
ns: not significant
NSG: Nod-scid gamma
OVA: ovalbumin (either full protein or the 257-264 amino acid sequence)
PBL: peripheral blood lymphocyte
PBS: phosphate-buffered saline
PD1: programmed death-1
PD-L1: programmed death ligand-1
PS: penicillin-streptomycin
RM-ANOVA: repeated measures analysis of variance
SEM: standard error of the mean
TCR: T cell receptor
TNFα: tumor necrosis factor-α
T-VEC: Talimogene laherparapvec
Y17: YUMMER1.7

