## Supplemental Figures for "The anti-virus T cell response dominates the anti-cancer response in oncolytic virus therapy"

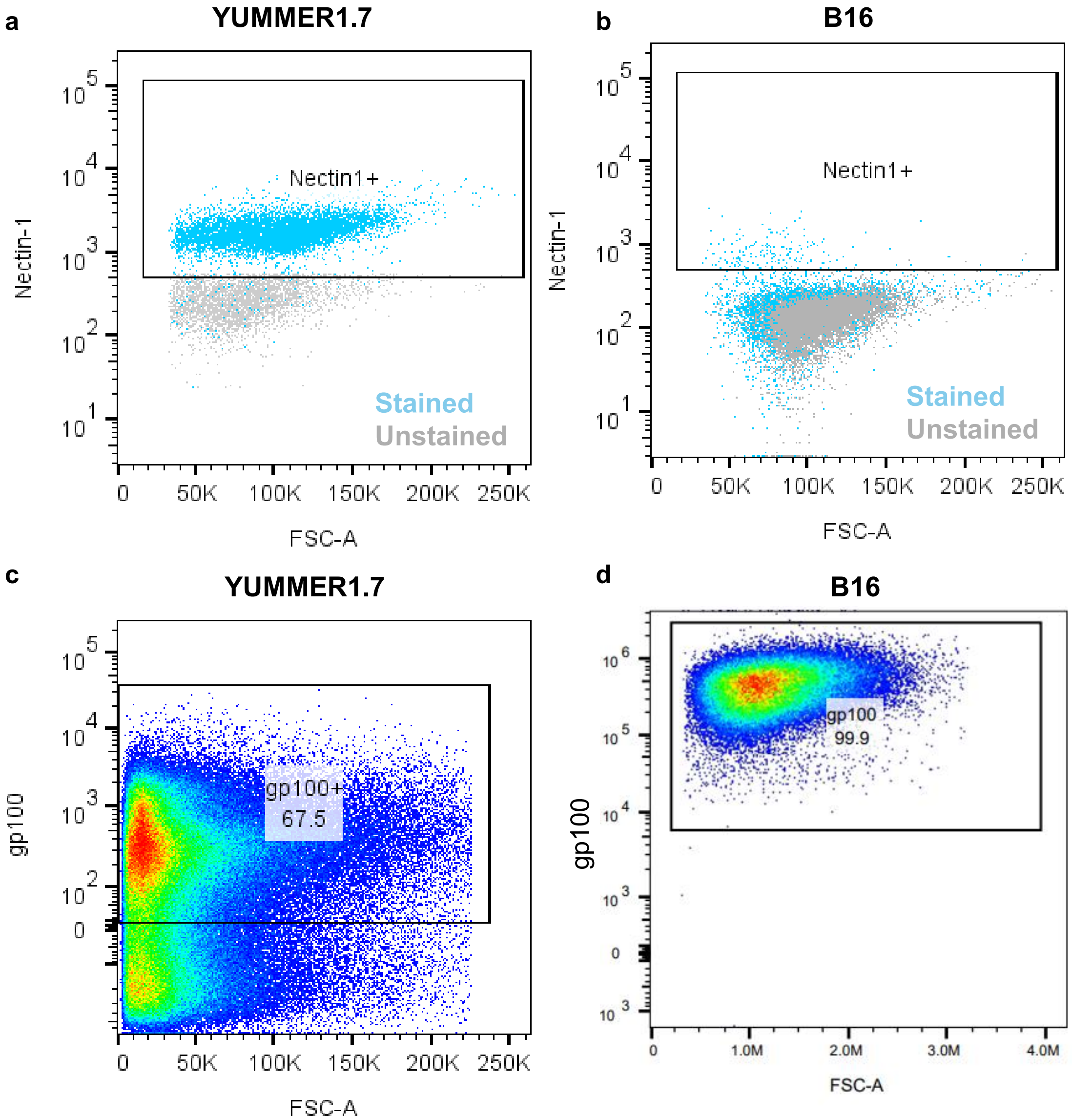
a
YUMMER1.7
Nectin1+
Stained
Unstained
Nectin-1
FSC-A
b
B16
Nectin1+
Stained
Unstained
Nectin-1
FSC-A
c
YUMMER1.7
gp100+
67.5
gp100
FSC-A
d
B16
gp100
99.9
gp100
FSC-A

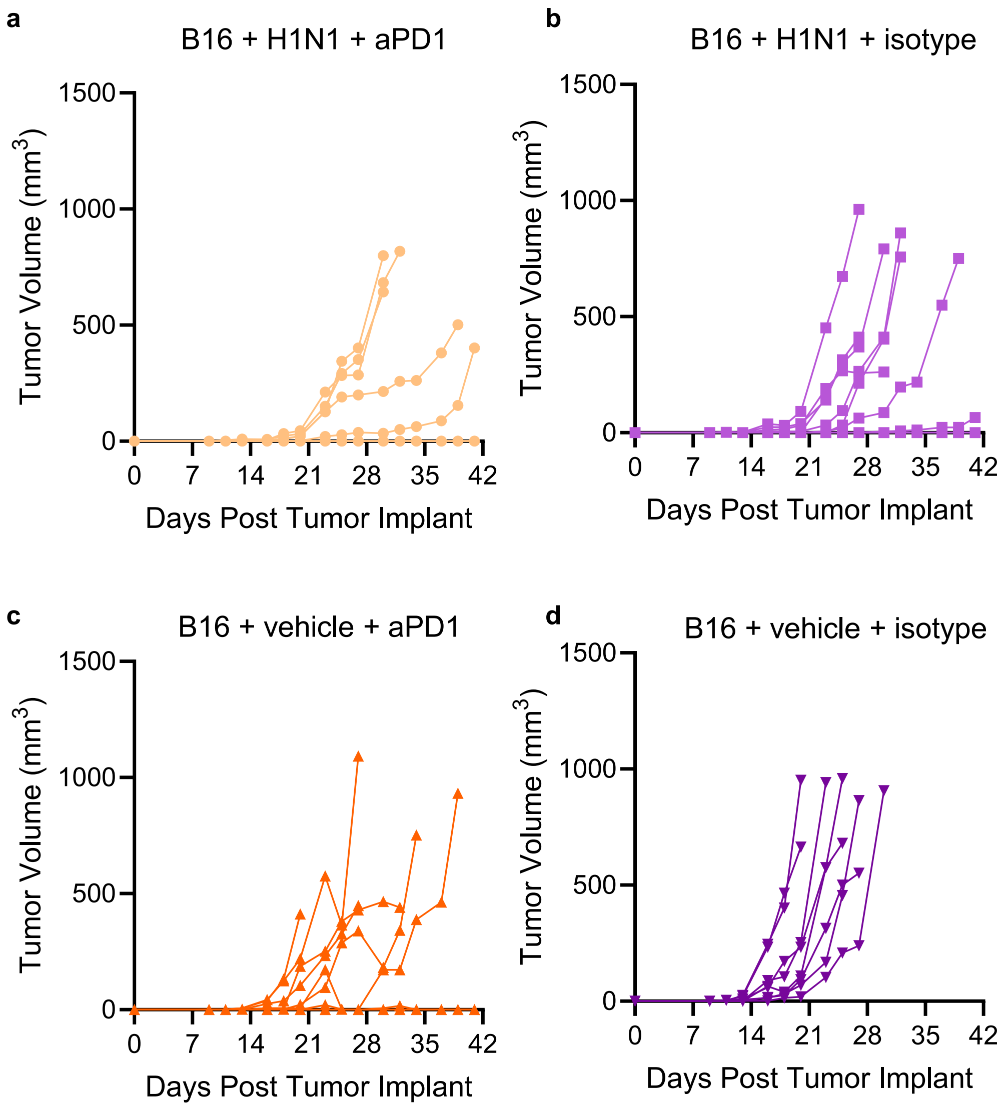

a
B16 + H1N1 + aPD1
b
B16 + H1N1 + isotype
c
B16 + vehicle + aPD1
d
B16 + vehicle + isotype
Tumor Volume (mm3)
Days Post Tumor Implant
0
7
14
21
28
35
42
500
1000
1500

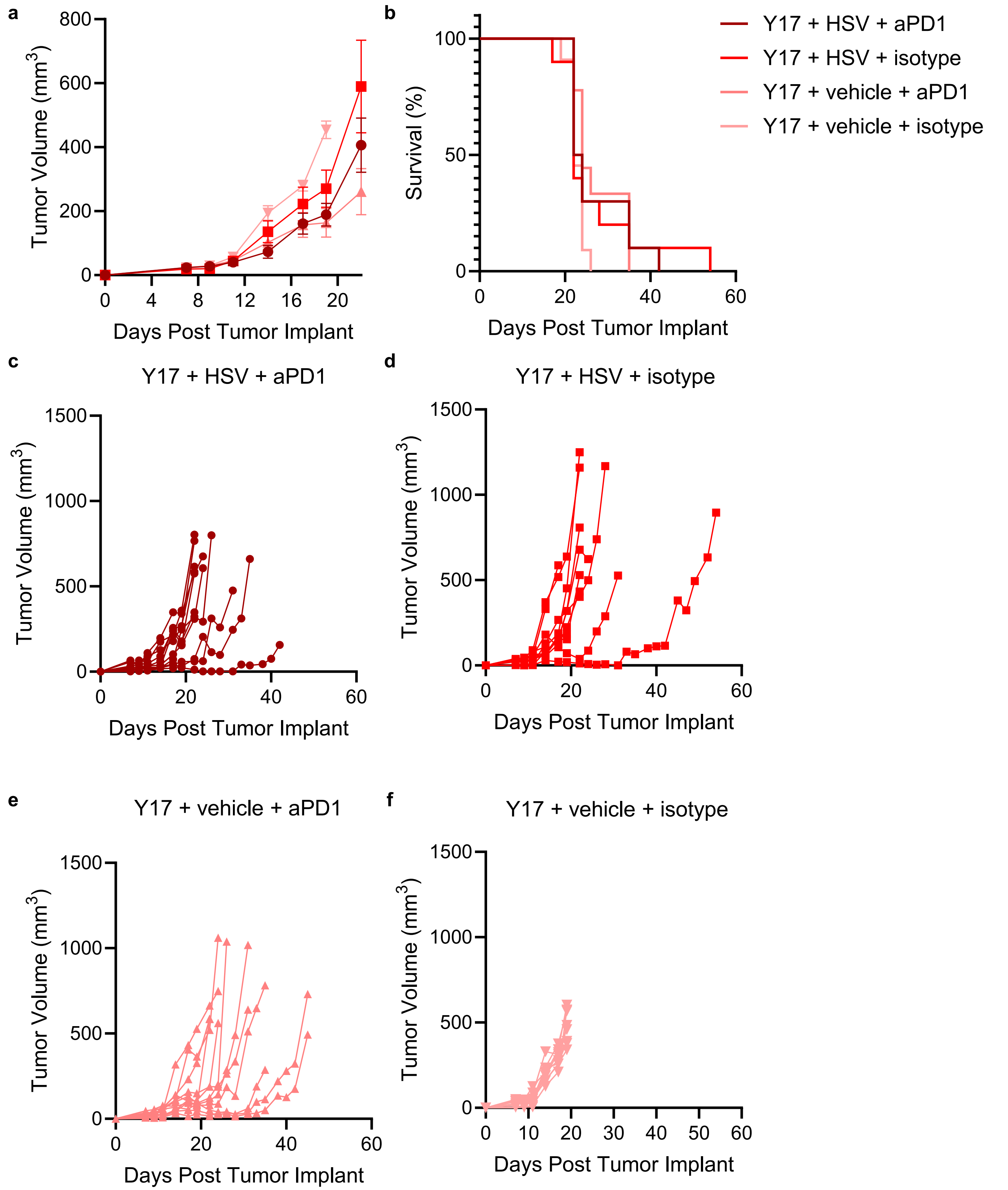

a
Tumor Volume (mm3)
Days Post Tumor Implant
b
Survival (%)
Days Post Tumor Implant
Y17 + HSV + aPD1
Y17 + HSV + isotype
Y17 + vehicle + aPD1
Y17 + vehicle + isotype
c
Y17 + HSV + aPD1
d
Y17 + HSV + isotype
e
Y17 + vehicle + aPD1
f
Y17 + vehicle + isotype

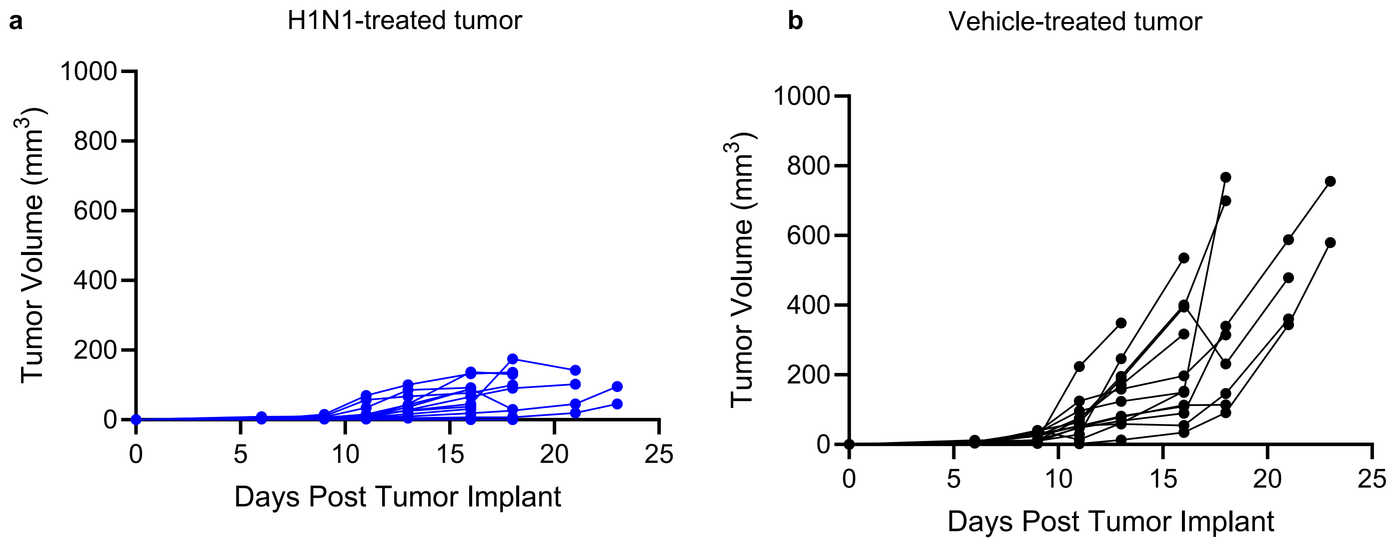
a
H1N1-treated tumor
Tumor Volume (mm3)
1000
800
600
400
200
0
0
5
10
15
20
25
Days Post Tumor Implant
b
Vehicle-treated tumor
Tumor Volume (mm3)
1000
800
600
400
200
0
0
5
10
15
20
25
Days Post Tumor Implant

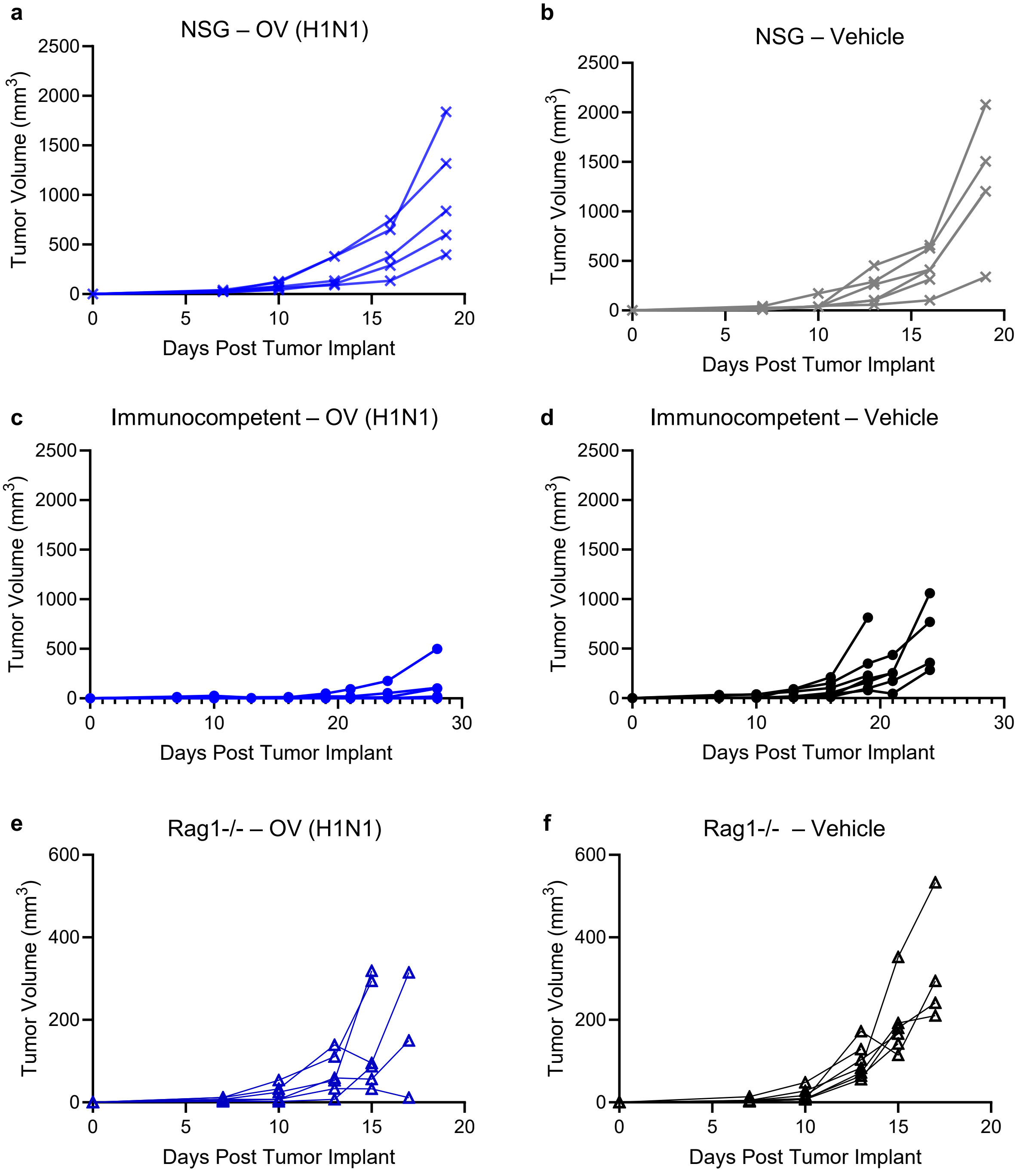
a
NSG – OV (H1N1)
Tumor Volume (mm3)
Days Post Tumor Implant
b
NSG – Vehicle
Tumor Volume (mm3)
Days Post Tumor Implant
c
Immunocompetent – OV (H1N1)
Tumor Volume (mm3)
Days Post Tumor Implant
d
Immunocompetent – Vehicle
Tumor Volume (mm3)
Days Post Tumor Implant
e
Rag1-/- – OV (H1N1)
Tumor Volume (mm3)
Days Post Tumor Implant
f
Rag1-/- – Vehicle
Tumor Volume (mm3)
Days Post Tumor Implant

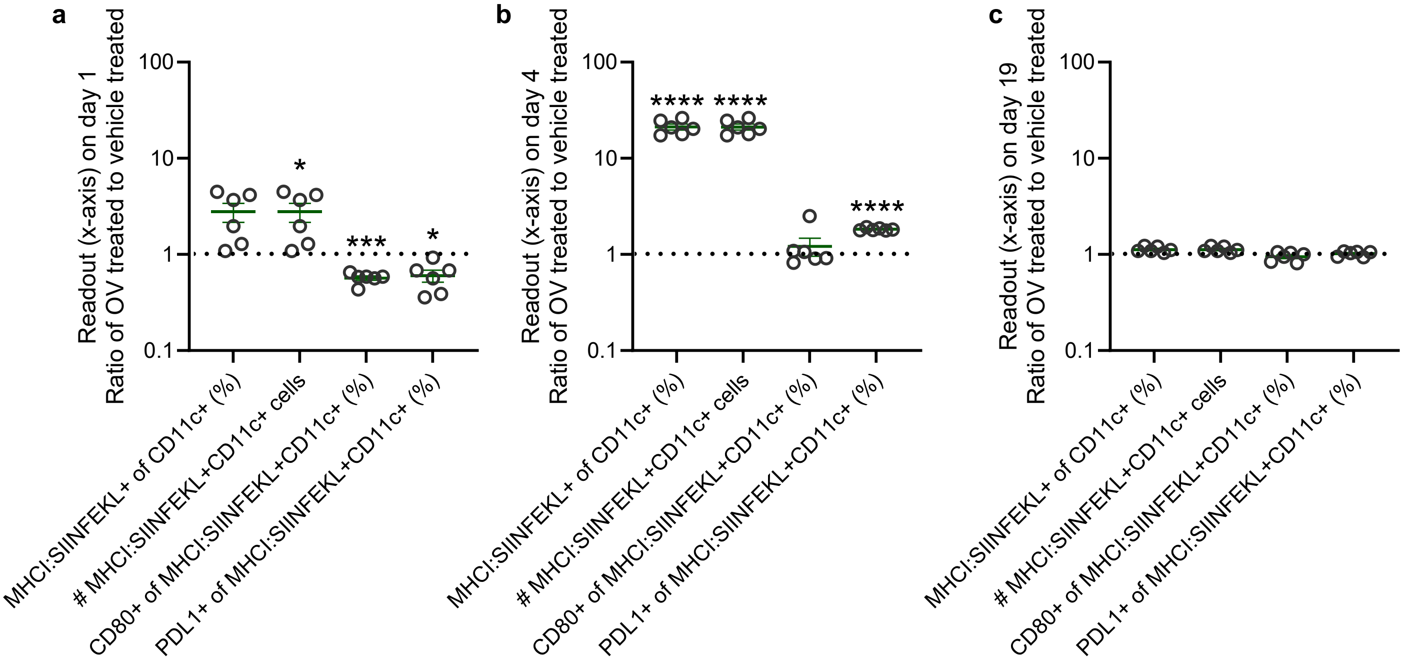
a
b
c
Readout (x-axis) on day 1
Ratio of OV treated to vehicle treated
Readout (x-axis) on day 4
Ratio of OV treated to vehicle treated
Readout (x-axis) on day 19
Ratio of OV treated to vehicle treated
100
10
1
0.1
*
***
****
MHCI:SIINFEKL+ of CD11c+ (%)
### MHCI:SIINFEKL+CD11c+ cells
CD80+ of MHCI:SIINFEKL+CD11c+ (%)
PDL1+ of MHCI:SIINFEKL+CD11c+ (%)

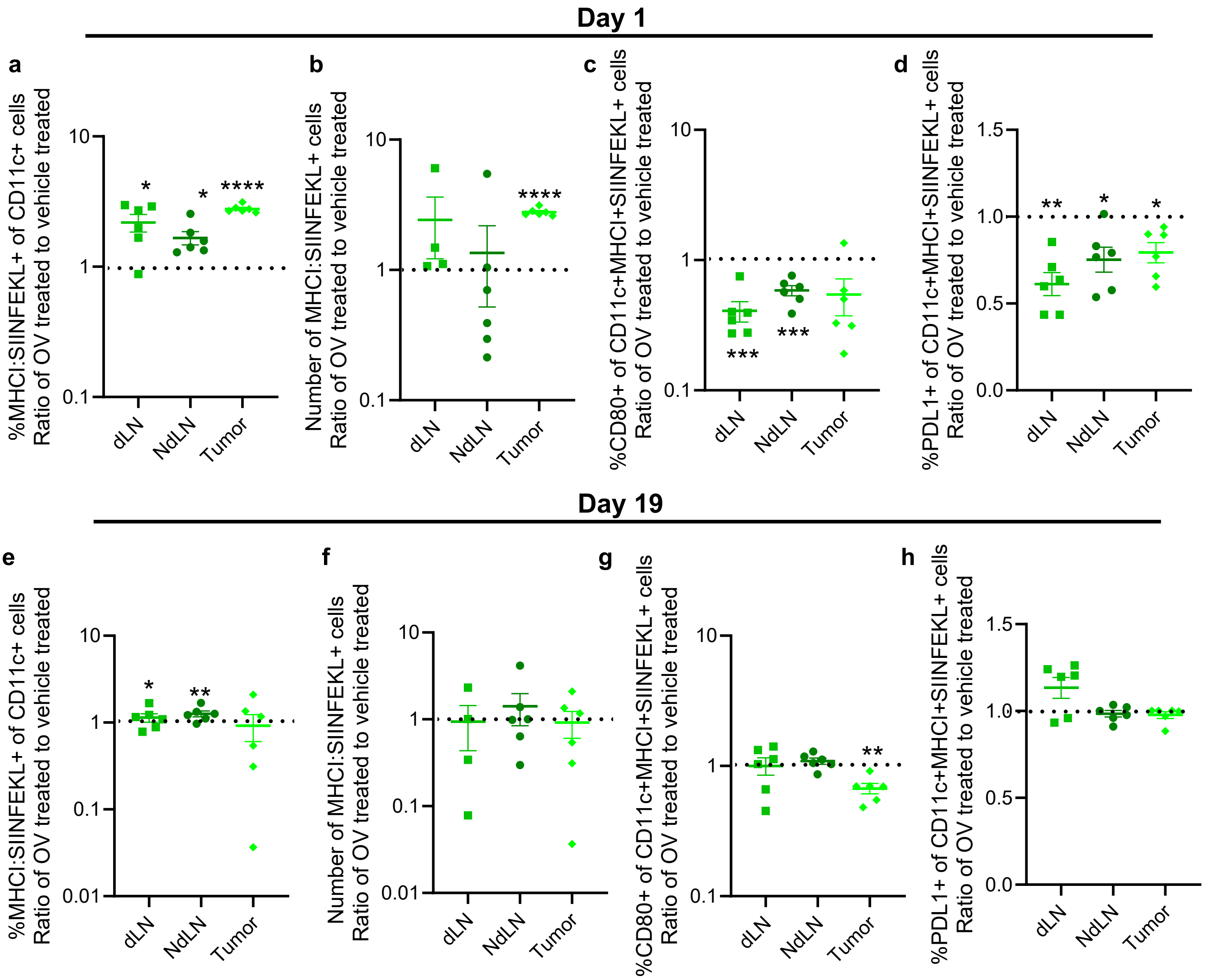
Day 1
a
%MHCI:SIINFEKL+ of CD11c+ cells
Ratio of OV treated to vehicle treated
10
1
0.1
*
*
****
dLN
NdLN
Tumor
b
Number of MHCI:SIINFEKL+ cells
Ratio of OV treated to vehicle treated
10
1
0.1
****
dLN
NdLN
Tumor
c
%CD80+ of CD11c+MHCI+SIINFEKL+ cells
Ratio of OV treated to vehicle treated
10
1
0.1
***
***
dLN
NdLN
Tumor
d
%PDL1+ of CD11c+MHCI+SIINFEKL+ cells
Ratio of OV treated to vehicle treated
1.5
1.0
0.5
0.0
**
*
*
dLN
NdLN
Tumor
Day 19
e
%MHCI:SIINFEKL+ of CD11c+ cells
Ratio of OV treated to vehicle treated
10
1
0.1
0.01
*
**
dLN
NdLN
Tumor
f
Number of MHCI:SIINFEKL+ cells
Ratio of OV treated to vehicle treated
10
1
0.1
0.01
dLN
NdLN
Tumor
g
%CD80+ of CD11c+MHCI+SIINFEKL+ cells
Ratio of OV treated to vehicle treated
10
1
0.1
**
dLN
NdLN
Tumor
h
%PDL1+ of CD11c+MHCI+SIINFEKL+ cells
Ratio of OV treated to vehicle treated
1.5
1.0
0.5
0.0
dLN
NdLN
Tumor

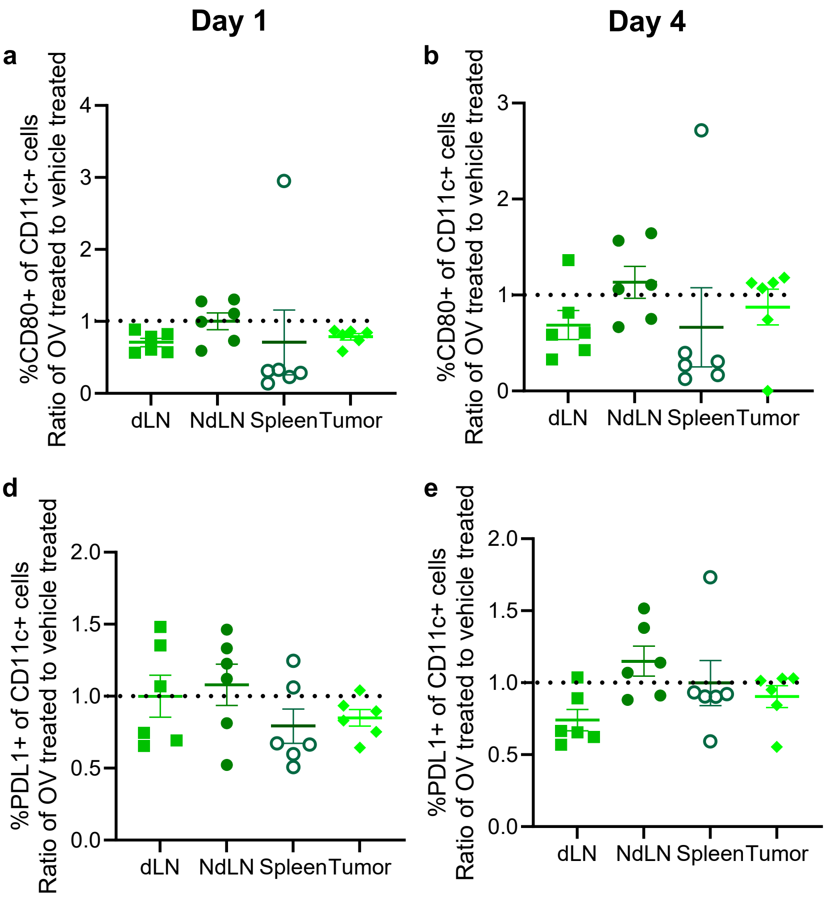

Day 1
Day 4
Day 19
a
b
c
d
e
f
%CD80+ of CD11c+ cells
Ratio of OV treated to vehicle treated
%PDL1+ of CD11c+ cells
Ratio of OV treated to vehicle treated
dLN
NdLN
Spleen
Tumor

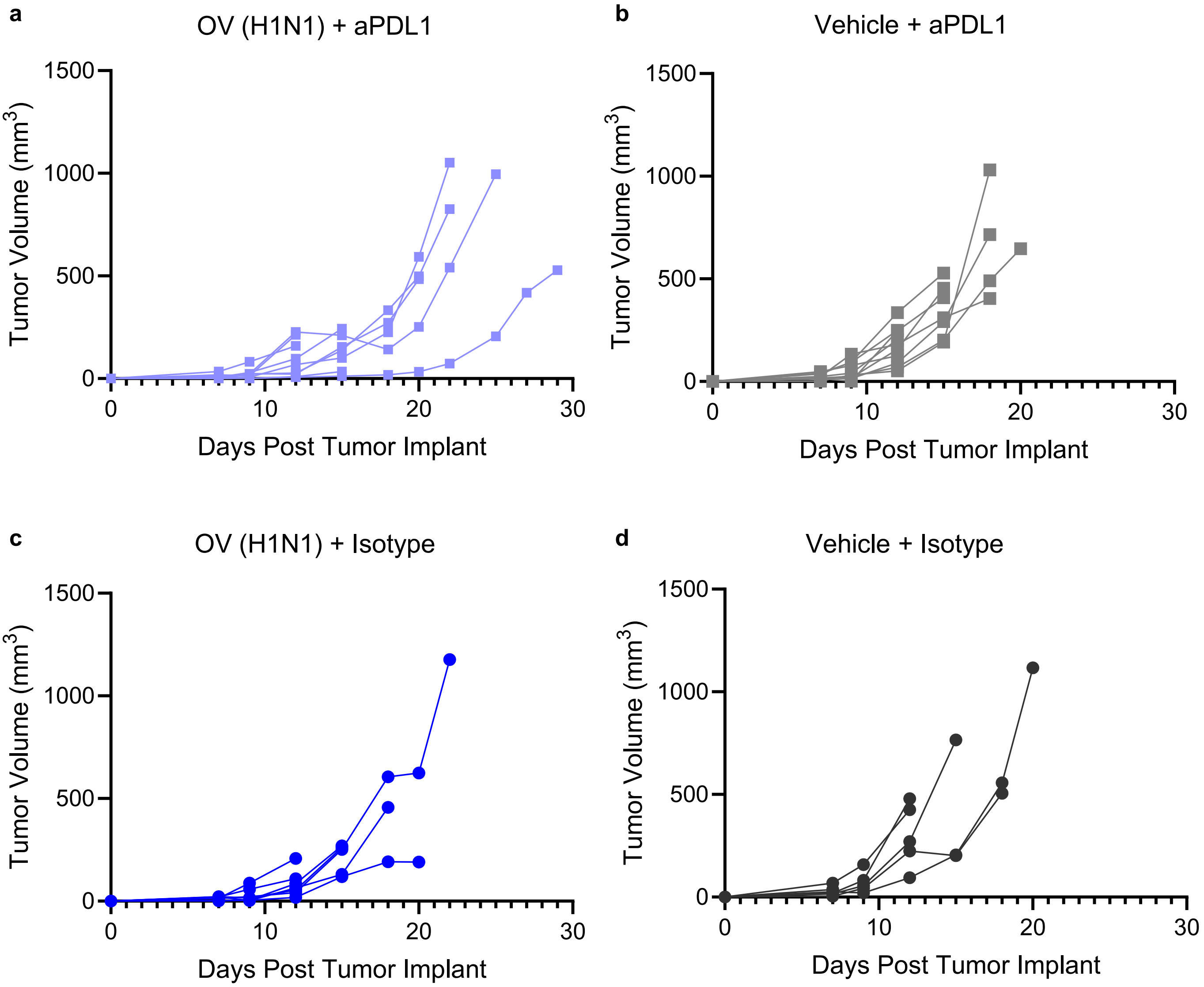
a
OV (H1N1) + aPDL1
b
Vehicle + aPDL1
c
OV (H1N1) + Isotype
d
Vehicle + Isotype
Tumor Volume (mm3)
Days Post Tumor Implant
0
500
1000
1500
0
10
20
30

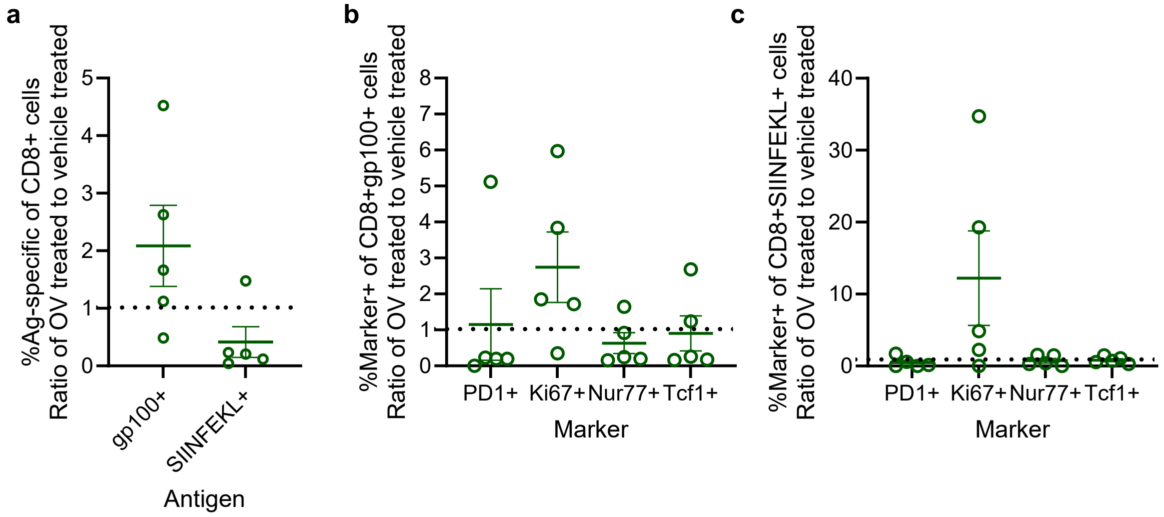

a
%Ag-specific of CD8+ cells
Ratio of OV treated to vehicle treated
gp100+
SIINFEKL+
Antigen
b
%Marker+ of CD8+gp100+ cells
Ratio of OV treated to vehicle treated
PD1+
Ki67+
Nur77+
Tcf1+
Marker
c
%Marker+ of CD8+SIINFEKL+ cells
Ratio of OV treated to vehicle treated
PD1+
Ki67+
Nur77+
Tcf1+
Marker

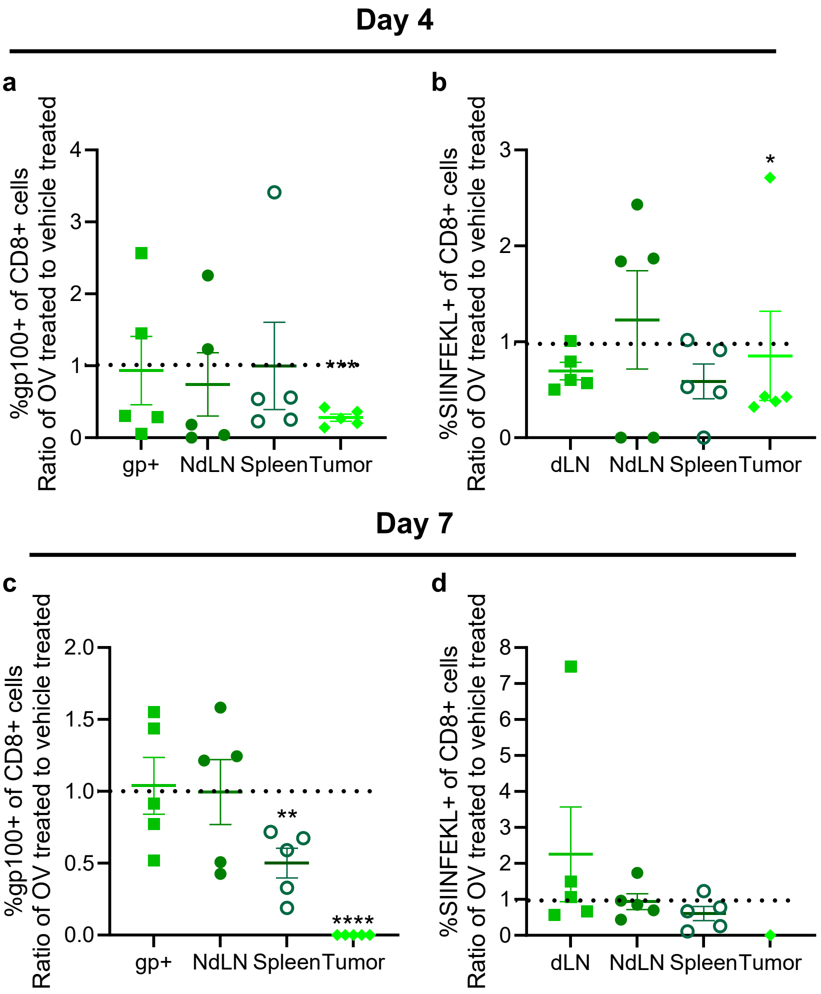
Day 4
a
%gp100+ of CD8+ cells
Ratio of OV treated to vehicle treated
gp+
NdLN
Spleen
Tumor
***
b
%SIINFEKL+ of CD8+ cells
Ratio of OV treated to vehicle treated
dLN
NdLN
Spleen
Tumor
*
Day 7
c
%gp100+ of CD8+ cells
Ratio of OV treated to vehicle treated
gp+
NdLN
Spleen
Tumor
**
****
d
%SIINFEKL+ of CD8+ cells
Ratio of OV treated to vehicle treated
dLN
NdLN
Spleen
Tumor

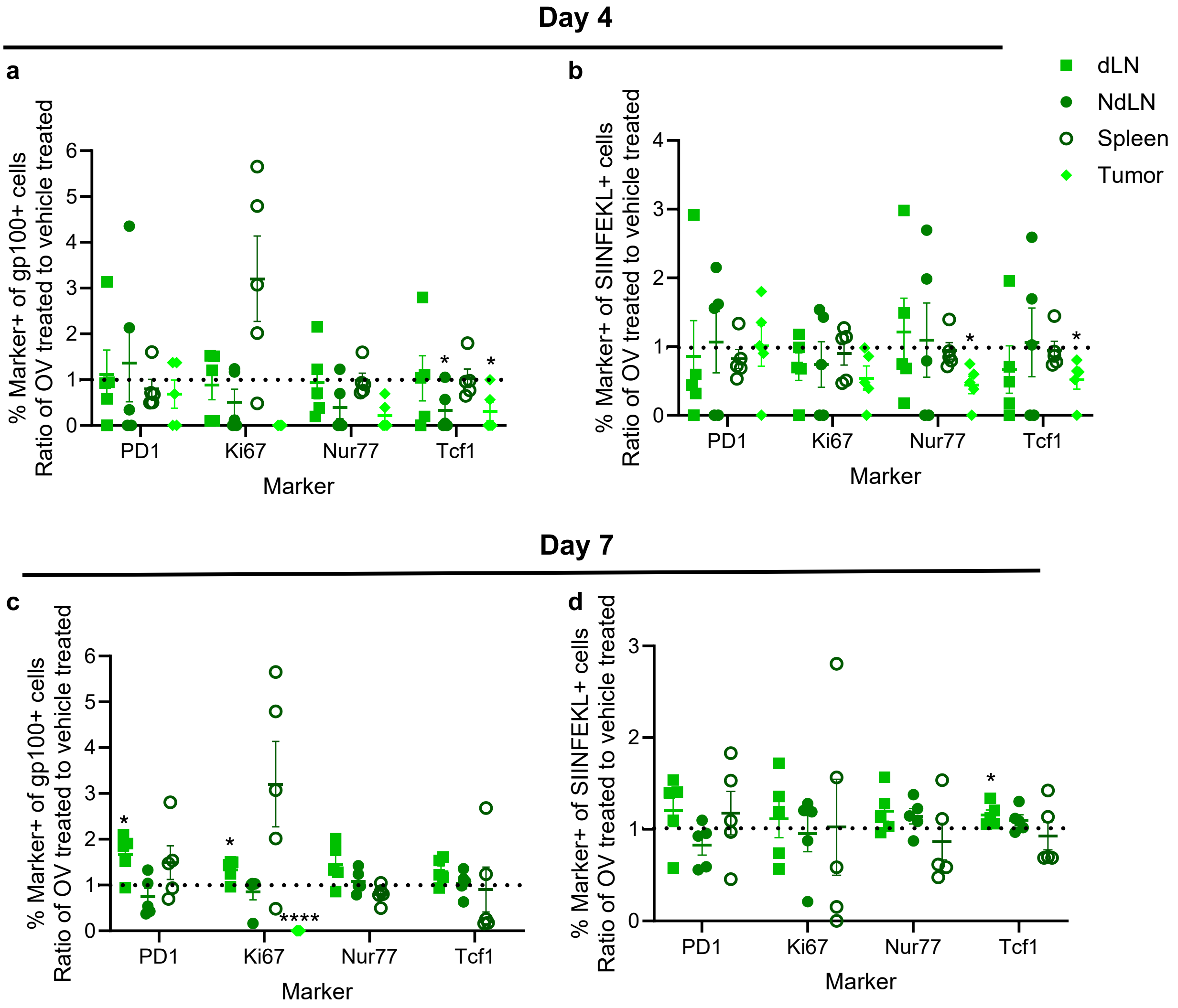

Day 4
a
% Marker+ of gp100+ cells
Ratio of OV treated to vehicle treated
0
1
2
3
4
5
6
PD1
Ki67
Nur77
Tcf1
Marker
*
*
b
% Marker+ of SIINFEKL+ cells
Ratio of OV treated to vehicle treated
0
1
2
3
4
PD1
Ki67
Nur77
Tcf1
Marker
*
*
dLN
NdLN
Spleen
Tumor
Day 7
c
% Marker+ of gp100+ cells
Ratio of OV treated to vehicle treated
0
1
2
3
4
5
6
PD1
Ki67
Nur77
Tcf1
Marker
*
*
****
d
% Marker+ of SIINFEKL+ cells
Ratio of OV treated to vehicle treated
0
1
2
3
PD1
Ki67
Nur77
Tcf1
Marker
*

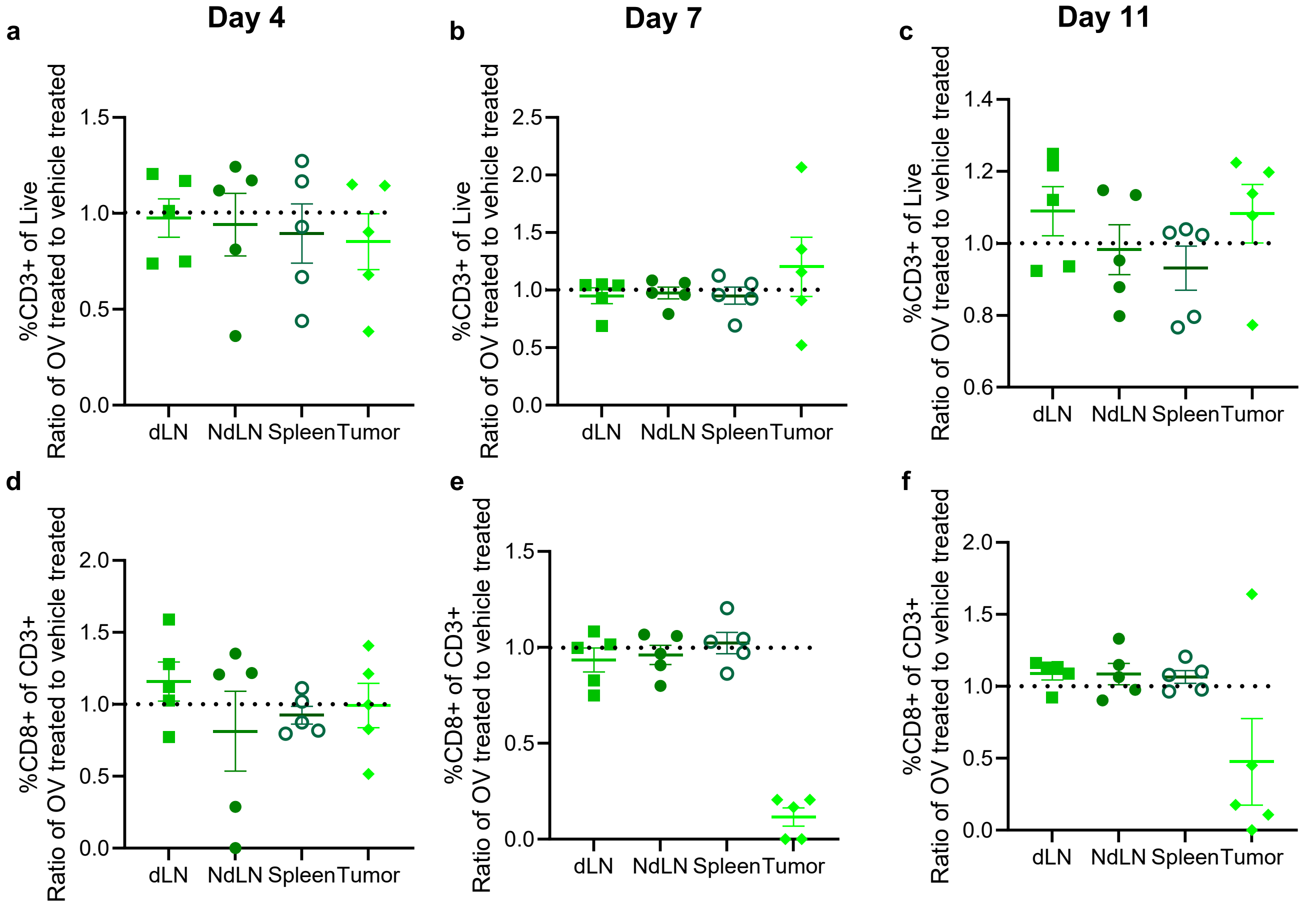
Day 4
Day 7
Day 11
a
b
c
d
e
f
%CD3+ of Live
Ratio of OV treated to vehicle treated
%CD8+ of CD3+
Ratio of OV treated to vehicle treated
dLN
NdLN
Spleen
Tumor

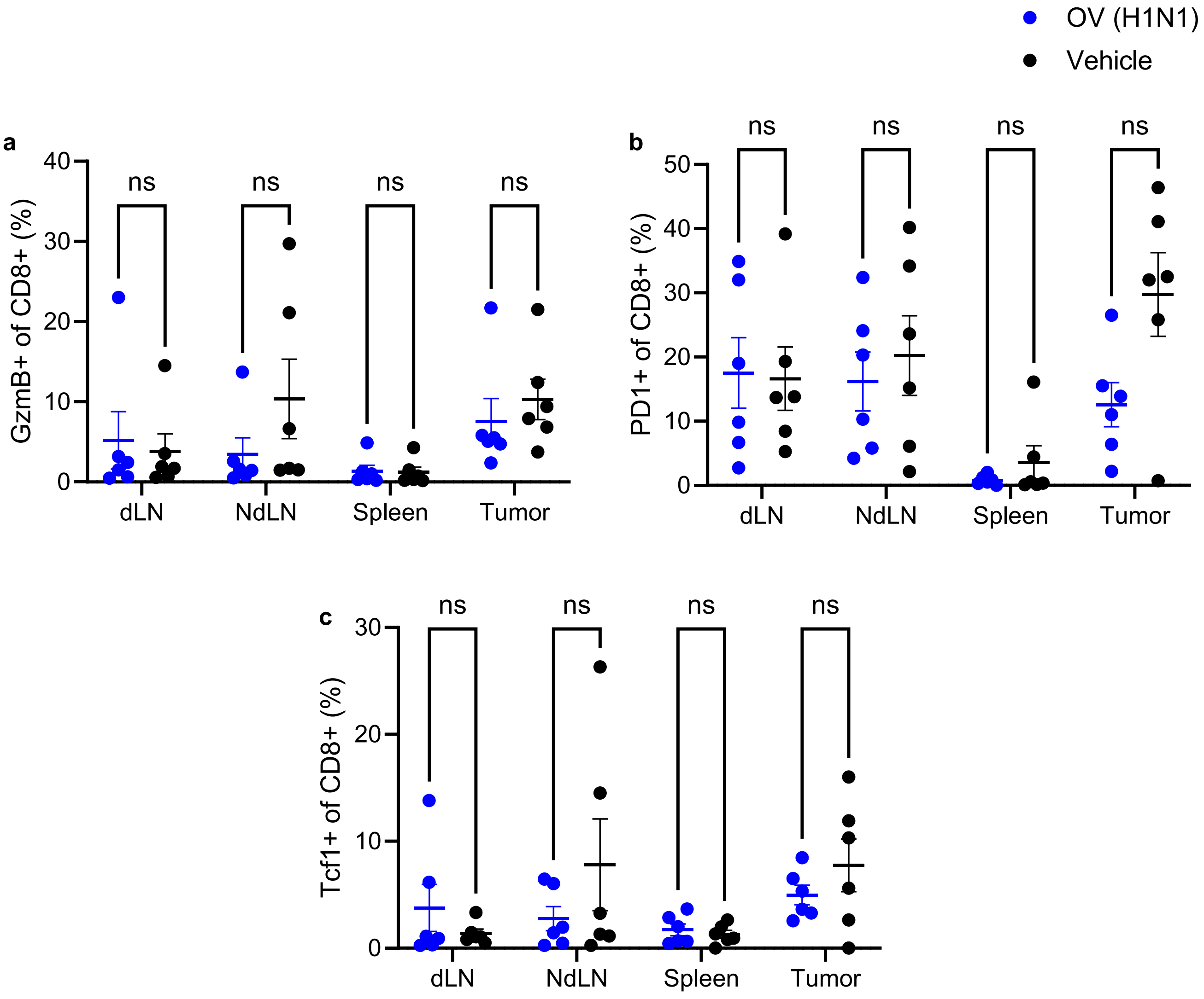

OV (H1N1)
Vehicle
a
GzmB+ of CD8+ (%)
b
PD1+ of CD8+ (%)
c
Tcf1+ of CD8+ (%)
ns
dLN
NdLN
Spleen
Tumor

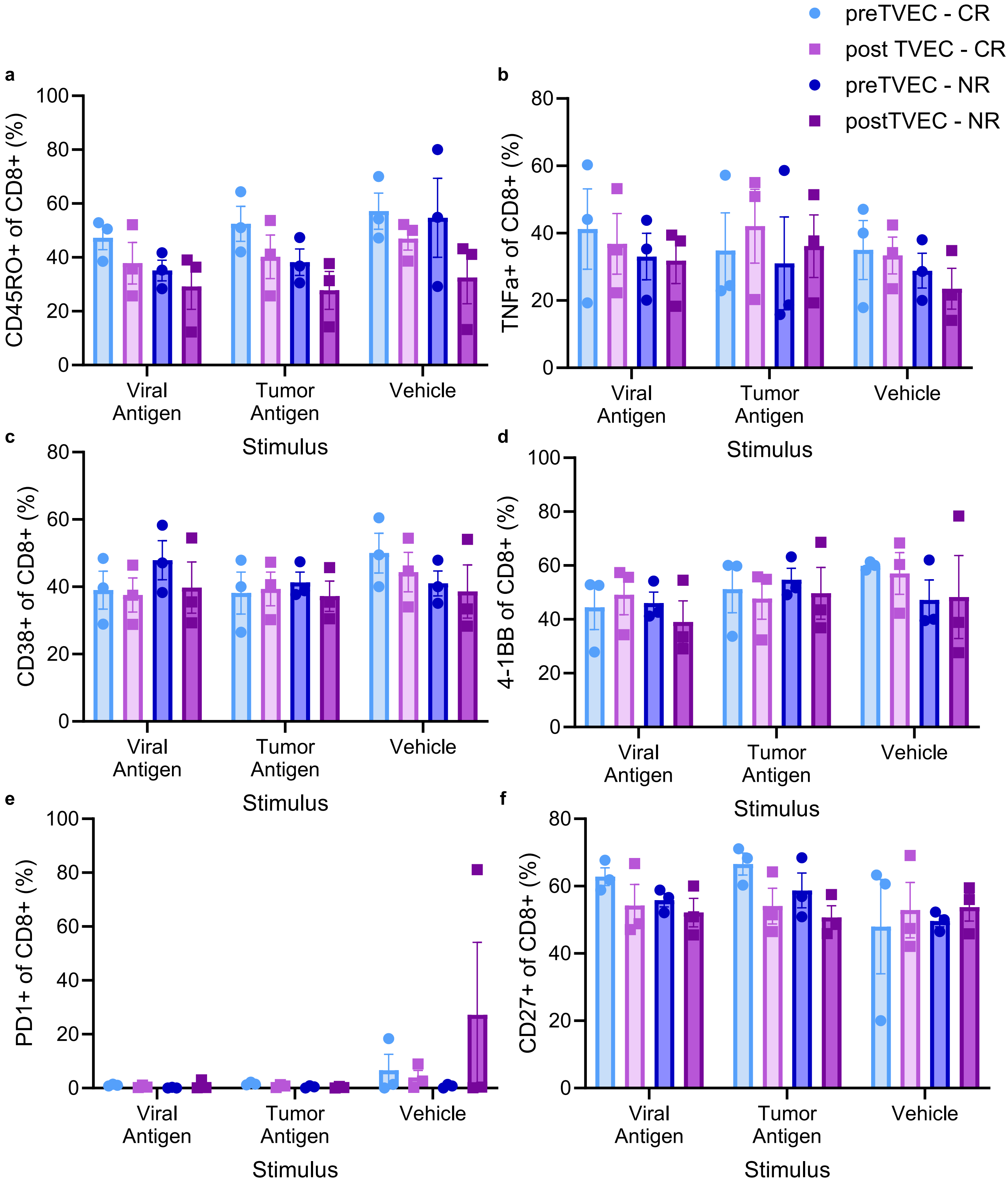

preTVEC - CR
post TVEC - CR
preTVEC - NR
postTVEC - NR
a
CD45RO+ of CD8+ (%)
b
TNFa+ of CD8+ (%)
c
CD38+ of CD8+ (%)
d
4-1BB of CD8+ (%)
e
PD1+ of CD8+ (%)
f
CD27+ of CD8+ (%)
Viral Antigen
Tumor Antigen
Vehicle
Stimulus
